# Type 1 lymphocytes accumulate in the thalamus and produce interferon-gamma to restrict seizure susceptibility after brain injury

**DOI:** 10.1101/2024.12.28.630606

**Authors:** Nicholas M. Mroz, Agnieszka Ciesielska, Nathan A. Ewing-Crystal, Leah C. Dorman, Rafael T. Han, Jerika J. Barron, Nicholas J. Silva, Audrey Magsig, Anna V. Molofsky, Ari B. Molofsky, Jeanne T. Paz

## Abstract

Chronic neural circuit hyperexcitability frequently emerges after brain injury, but endogenous mechanisms constraining runaway activity remain poorly understood. Here, we reveal that the adaptive immune system acts as a homeostatic brake on network excitability following traumatic brain injury (TBI). In mice, cortical trauma triggered a delayed infiltration of interferon-γ (IFNγ)–producing type 1 lymphocytes into the sensory thalamus. Rather than driving pathology, IFNγ signaling directly in neurons restricted thalamocortical network hyperexcitability. This protective axis was tonically regulated; depleting CD4⁺ T cells de-repressed local non-CD4⁺ type 1 lymphocytes, elevating IFNγ signaling and protecting from seizures. A single dose of exogenous IFNγ abolished hypersynchronous circuit bursting and rescued injury-induced seizure incidence, severity, and mortality, establishing a therapeutic framework for safeguarding circuit stability after brain injury.

## Introduction

Traumatic brain injury (TBI) is a leading cause of worldwide morbidity, triggering a critical latent period of secondary pathogenesis during which functionally intact neural circuits transform into hypersynchronous, seizure-prone networks (*1–3*). Because these early manifestations of circuit instability are highly predictive of chronic neurological sequelae, disrupting this pathogenic trajectory during the latent window is essential for disease prevention (*4*). Chronic neuroinflammation is widely implicated as a primary driver of this maladaptive circuit reorganization (*4*); however, clinical efforts to halt progression using non-specific anti-inflammatory drugs have systematically failed (*5*, *6*). This therapeutic failure underscores that the post-injury inflammatory cascade is not uniformly destructive. Disentangling these complex responses to identify and ultimately boost endogenous, adaptive immune factors that actively restrict circuit hyperexcitability and enforce network homeostasis represents a critical frontier for disease-modifying intervention (*7–9*).

Reciprocal thalamo-cortico-thalamic neuronal networks are vulnerable to this post-injury transformation and represent a primary locus of heightened seizure risk (*10–12*). Following primary cortical trauma, secondary damage cascades into the physically distant but functionally connected thalamus as cortical-projecting thalamic neurons degenerate retrogradely, precipitating chronic thalamic gliosis and neuroinflammation (*13–15*). Secondary thalamic damage can drive pro-epileptogenic circuit alterations dependent on innate immune activation and complement-mediated neurodegeneration (*10*, *12*, *16*). Because secondary thalamic neuroinflammation is a universal feature of cortical injuries independent of primary etiology (i.e., stroke, TBI) (*10*, *13*, *17*), this distinct spatial compartment provides an ideal microenvironment to isolate whether the chronic inflammatory niche harbors undiscovered adaptive immune components attempting to stabilize a collapsing circuit.

Neuroinflammation includes highly distinct immune cell types, spatial architectures, and central nervous system (CNS) targets. Canonically, TBI drives early recruitment of innate immune cells that cooperate with local microglia, followed by a delayed recruitment of T cells into the brain parenchyma (*4*, *18*). While these CNS-resident lymphocytes accumulate after many CNS perturbations (*18–30*), how they actively dialogue with parenchymal cells during the post-injury latent window to impact downstream neurological sequelae and modulate network excitability remains unknown.

Here, a broad immunophenotyping screen across multiple brain regions revealed a distinct population of type 1 lymphocytes that selectively and robustly infiltrated the sensory thalamus weeks after cortical TBI. By combining high-dimensional flow cytometry and confocal imaging, we discovered that these lymphocytes produce interferon-γ (IFNγ), which elicits an IFN-response within the sensory thalamus, including in excitatory thalamocortical neurons. Using electrophysiology, seizure risk assays, and genetic tools, we identified IFNγ signaling, particularly within neurons, to be an essential homeostatic brake against post-TBI circuit hyperexcitability. Finally, we show that systemic administration of exogenous IFNγ during this critical post-injury latent window is sufficient to normalize hyper-synchronous thalamocortical activity and rescue seizure susceptibility, establishing an unexpected therapeutic strategy to mitigate secondary neurological sequelae.

## Results

### A broad immune screen reveals delayed, anatomically constrained type 1 lymphocyte infiltration within the sensory thalamus after TBI

Cortical TBI leads to secondary and persistent neuroinflammation in the physically distant but functionally connected ipsilateral thalamus (Fig. 1A) (*10*, *17*). To systematically define the cellular composition and temporal kinetics of the immune response in this secondary niche, we performed flow cytometric profiling of the perilesional cortex, ipsilateral thalamus, and ipsilateral hippocampus at sequential intervals post-injury (fig. S1A). We identified a population of lymphocytes that infiltrated the ipsilateral thalamus with delayed kinetics as compared with the perilesional cortex (Fig. 1B). Whereas cortical lymphocyte numbers peaked at ∼14 days post injury (dpi), thalamic infiltration manifested as a delayed wave that peaked at 30–40 dpi and remained chronically elevated through at least ∼80 dpi (Figs. 1B and C). These infiltrates consisted primarily of T cells and natural killer (NK) cells, with few accompanying B cells (Fig. 1, D to G; fig. S1, B and C). Across both regions, lymphocytes were predominately CD8^+^ T cells, conventional CD4^+^ T cells (T_conv_, CD4^+^ FoxP3^-^), and NK cells (NK1.1^+^), sharing synchronized infiltration kinetics (Fig. 1, D to G, fig. S1, B and C).

**Fig. 1.**
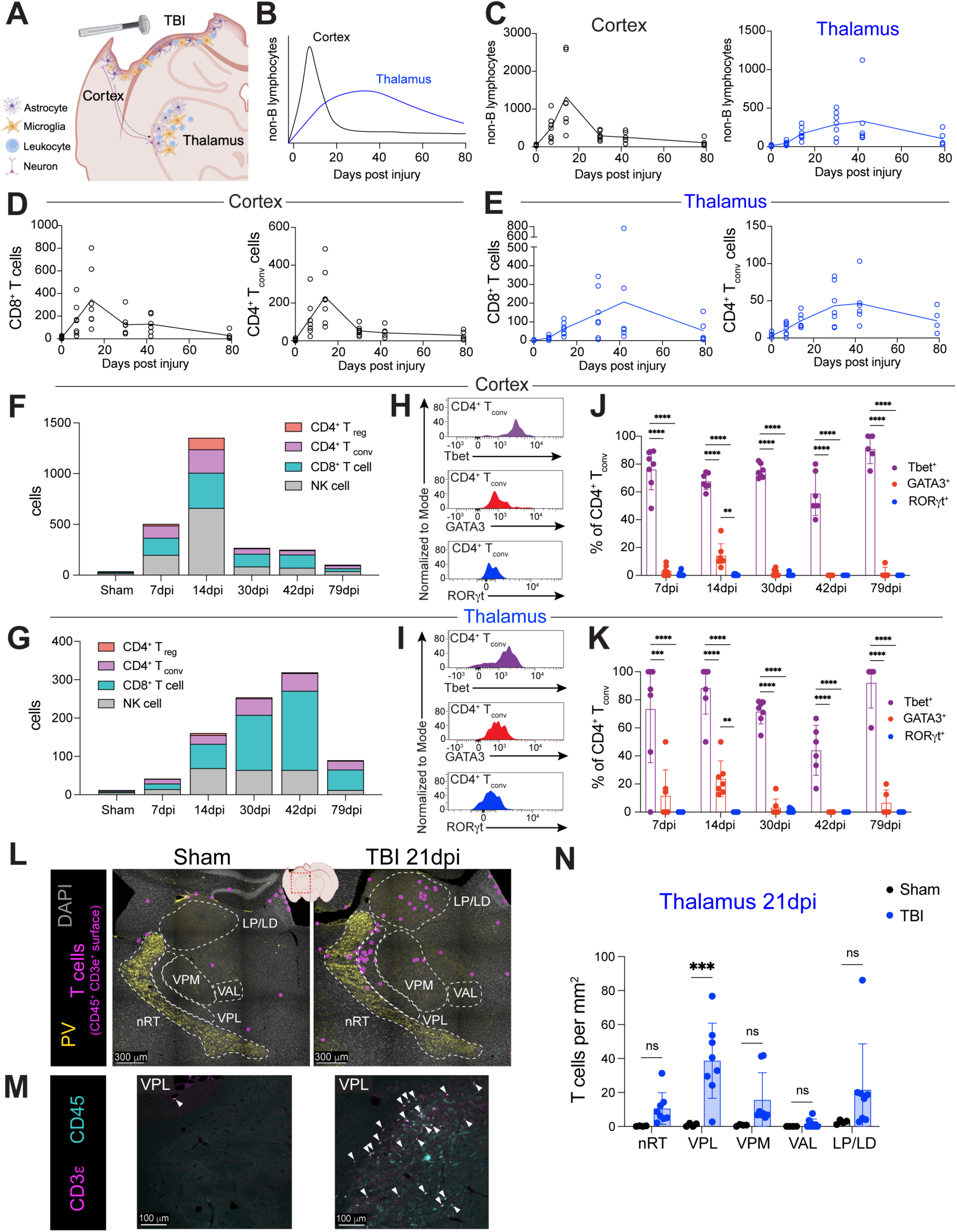
Type 1 adaptive lymphocytes infiltrate and persist within the thalamus following TBI. **(A)** Experimental schematic of the controlled cortical impact model of TBI at 21 days post-injury (dpi), illustrating localized perilesional cortical and retrograde thalamic gliosis (astrocytes and microglia) alongside peripheral leukocyte infiltration across a coronal murine brain section. **(B)** Schematized kinetic profiles tracking the spatiotemporal infiltration of non-B lymphocytes into the perilesional cortex and ipsilateral thalamus over time. **(C)** Absolute quantification of infiltrating non-B lymphocytes (CD45^hi^ CD11b^−^ CD19^−^ Thy1.2^+^) isolated from the perilesional cortex and ipsilateral thalamus across a longitudinal post-TBI timecourse (see fig. S1A for gating strategy). p = 0.0228 (cortex) and p = 0.0122 (thalamus). **(D and E)** Temporal dynamics showing absolute numbers of CD8⁺ T cells and conventional CD4^+^ FoxP3^−^ T cells (T_conv_) in the perilesional cortex **(D)** and ipsilateral thalamus **(E)**. **(F and G)** Stacked kinetic distribution of dominant lymphocytes, including NK/ILC1 (NK1.1^+^), CD8^+^ T, CD4^+^ FoxP3^−^ T_conv_, and CD4^+^ FoxP3^+^ T_reg_ cells, within the perilesional cortex (F) or ipsilateral thalamus (G). **(H and I)** Representative flow cytometric histograms showing intracellular expression of lineage-defining transcription factors (Tbet, GATA3, and RORγt) in (H) cortical CD4^+^ T_conv_ cells at 7 dpi or (I) thalamic CD4^+^ T_conv_ cells at 30 dpi. **(J and K)** Relative frequencies of Tbet^+^, GATA3^+^, and RORγt^+^ transcription factor expression within CD4^+^ T_conv_ cells isolated from perilesional cortex (J) or ipsilateral thalamus (K). **(L)** Representative confocal images of the ipsilateral thalamus in sham and TBI mice (21 dpi) showing parenchymal T cell infiltration (magenta circles; 3D surface-rendered CD45^+^ CD3ε^+^ reconstructions) relative to anatomical subregions (white dashed lines). Parvalbumin immunostaining (PV) delineates boundaries of the reticular thalamic nucleus (nRT). **(M)** Representative confocal images of VPL showing native CD45 and CD3å stains from ipsilateral thalamus of sham and TBI mice (21 dpi). Arrowheads indicate CD45^+^ CD3ε^+^ T cells **(N)** Spatial density of T cells (CD45^+^ CD3ε^+^) normalized per unit area (mm^2^) across distinct anatomical thalamic nuclei in sham versus TBI cohorts at 21 dpi. **Data and Statistics:** Main panels represent mean ± SD; panels (F) and (I) depict mean values only. Individual data points represent unique biological replicates (independent mice). Thalamic subregions: reticular nucleus (nRT), ventral posterolateral (VPL), ventral posteromedial (VPM), ventral anterolateral (VAL), and lateral posterior/lateral dorsal (LP/LD). ● **Sample sizes:** For longitudinal timecourses (C to K), n = 7 mice/group for sham (0 dpi), 7, 14, and 30 dpi; n = 6 for 42 dpi; and n = 5 for 79 dpi. For confocal imaging (N), sham n = 4, TBI n = 8. ● **Statistics:** Evaluated via mixed-effects model (REML) for (B to E): For (D), p = 0.0016 (CD8^+^ T cells) and p < 0.0001 (CD4^+^ T_conv_). For (E), p = 0.0482 (CD8^+^ T cells) and p = 0.0007 (CD4^+^ T_conv_). Evaluated via one-way ANOVA with Tukey’s multiple-comparisons test within individual timepoints for (H) and (K), and via two-way ANOVA with Šidák’s multiple-comparisons test for (N). ns = not significant; *p < 0.05; **p < 0.01; ***p < 0.001; ****p < 0.0001.

We next interrogated the types of brain-infiltrating lymphocytes after TBI. Both cortical and thalamic CD8^+^ T cells and NK cells expressed the canonical type 1 transcription factor Tbet (fig. S1, D and E), as expected (*31*). CD4^+^ T_conv_ cells across both of these brain regions also preferentially expressed Tbet over the type 2 (GATA3) or type 3/17 (RORγt) master transcriptional regulators, demonstrating a highly uniform type 1 polarization within the post-injury brain (Fig. 1, H to K; fig. S1, D and E) (*32*, *33*). Across timepoints, most thalamic T cells expressed a core phenotypic signature of brain tissue-resident memory (TRM) cells, characterized by CD44 and CD69 expression and lack of the lymph node-homing receptor CD62L (fig. S1, F to H) (*19*, *20*, *22*, *23*, *29*, *34*). Some of these cells co-expressed the TRM marker CD103 (Integrin α_E_), which was previously identified on a subset of brain-resident T cells (*19*, *20*, *22*, *23*, *29*), and the brain TRM signature was distinct from circulating blood lymphocytes (fig. S1, F to H). Lymphocyte infiltration into the ipsilateral hippocampus was negligible (fig. S1, I and J), establishing that post-traumatic lymphocyte infiltration is anatomically partitioned. These data indicate that the post-traumatic thalamus supports a specialized, tissue-resident type 1 immune microenvironment poised for interferon-γ (IFNγ) production.

Because secondary thalamic pathogenesis is anatomically restricted (*10*, *17*), we mapped the precise spatial organization of these infiltrating T cells using confocal microscopy. At 21 dpi, T cells (CD45^+^ CD3ε^+^) selectively accumulated within specific thalamic subregions functionally wired to the primary somatosensory cortical lesion, most notably in the glutamatergic ventroposterolateral (VPL) nucleus (Fig. 1, L to N). In contrast, T cells were not elevated in the ventroanterolateral (VAL) motor thalamus, despite its close anatomical proximity to the sensory thalamus (Fig. 1N). Together, these findings reveal that cortical TBI recruits a delayed wave of T cells that map precisely onto secondarily injured somatosensory thalamic nuclei.

### Infiltrating type 1 lymphocytes produce IFNγ within the injured sensory thalamus

Because type 1 lymphocyte polarization is functionally defined by the secretion of IFNγ (*31*, *32*), we quantified cytokine dynamics within microdissected brain regions across a subacute-to-chronic post-injury timeline. Quantitative immunoassays revealed a progressive, localized accumulation of IFNγ that was more concentrated within the ipsilateral thalamus than either the perilesional cortex or the hippocampus (Fig. 2, A to D). This inflammatory signature was pathway-specific: other canonical lymphocyte-derived cytokines, including interleukin-2 (IL-2) and the type 2 cytokine IL-13, were unaltered (fig. S2, A to C).

**Fig. 2.**
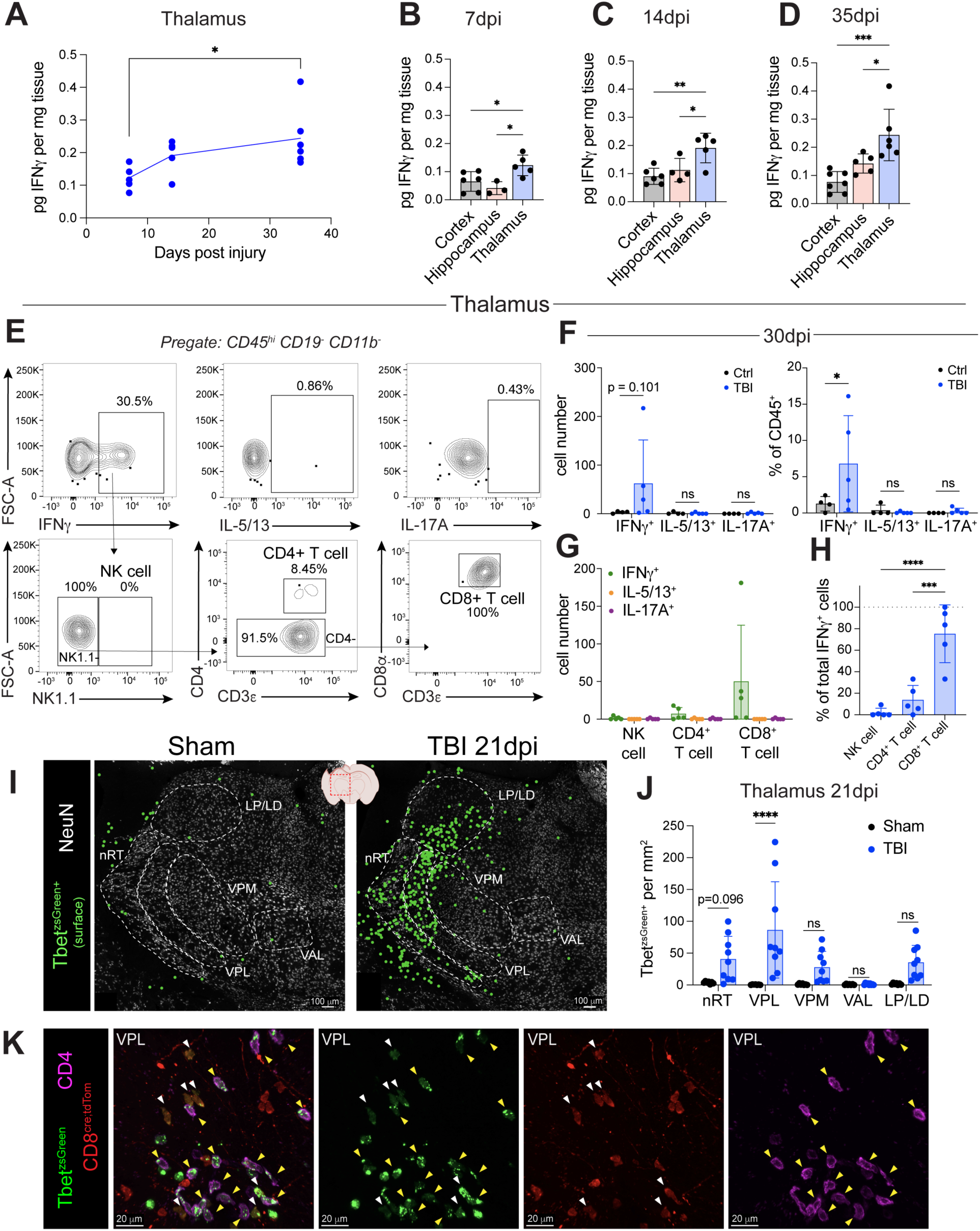
IFNγ is elevated in the thalamus following TBI and is produced primarily by infiltrating CD8^+^ T cells. **(A)** Temporal profile of IFNγ protein expression within the ipsilateral thalamus at 7, 14, and 35 days post-injury (dpi) measured via multiplex bead immunoassay on micro-dissected tissue homogenates. **(B to D)** Spatial tracking of IFNγ concentrations compared across the perilesional cortex, ipsilateral hippocampus, and ipsilateral thalamus at 7 dpi **(B)**, 14 dpi **(C)**, and 35 dpi **(D)**. All cytokine values are normalized to total tissue weight (mg). **(E)** Representative flow cytometric plots displaying intracellular expression of IFNγ, IL-5/13, and IL-17A within stimulated thalamic leukocytes (CD45^hi^ CD11b^-^ CD19^-^ cells) at 41 dpi (top row). Secondary gating of the total IFNγ^+^ population (bottom row) delineates the relative abundance of NK/ILC1s (NK1.1^+^), CD4^+^ T cells, and CD8^+^ T cells. **(F)** Total cell numbers (left) and relative frequencies within total CD45^+^ brain-infiltrating leukocytes (right) of IFNγ^+^, IL-5/13^+^, and IL-17A^+^ cells in the thalamus of sham versus TBI mice at 30 dpi. **(G)** Absolute quantification of cytokine-producing (IFNã^+^, IL-5/13^+^, IL-17A^+^) lymphoid subsets (NK/ILC1, CD4^+^ T, and CD8^+^ T cells) in the injured thalamus at 30 dpi. **(H)** Immunophenotypic composition of the total thalamic IFNã^+^ pool at 41 dpi, showing the relative frequency of NK/ILC1s, CD4^+^ T cells, and CD8^+^ T cells. **(I)** Representative confocal images of the ipsilateral thalamus from Tbet reporter mice (*Tbx21^zsGreen^*) at 21 days after sham or TBI surgeries, displaying Tbet-expressing cells (green circles; 3D surface reconstructions) and anatomical subregions (white dashed lines) overlaid on NeuN stain. **(J)** Spatial density of reporter-positive (Tbet^zsGreen+^) cells normalized per unit area (mm^2^) across distinct anatomical thalamic subregions in sham versus TBI cohorts at 21 dpi. **(K)** High-power confocal image of the ventroposterolateral (VPL) thalamic nucleus from a triple-transgenic reporter mouse (*Tbx21^zsGreen^; Cd8a^E8i-cre/+^; Rosa26^Ai14/+^*) at 21 dpi. Images display merged (left) and individual single-channel fluorescence for native Tbet^zsGreen^, CD8^cre;tdTomato^, and CD4 stains. White arrowheads indicate Tbet-expressing cytotoxic T cells (Tbet^zsGreen+^ CD8^cre;tdTom+^); yellow arrowheads mark Tbet-expressing helper T cells (Tbet^zsGreen+^ CD4^+^). **Data and Statistics:** Bar graphs and data points represent mean ± SD from individual mice (biological replicates). ● **Sample Sizes (n):** For thalamic cytokine kinetics (A), 7 dpi n = 5, 14 dpi n = 5, 35 dpi n = 6. For regional cytokine mapping (B to D), 7 dpi (cortex n = 6, hippocampus n = 3, thalamus n = 5); 14 dpi (cortex n = 6, hippocampus n = 4, thalamus n = 5); 35 dpi (cortex n = 7, hippocampus n = 5, thalamus n = 6). For flow cytometry (F to H), sham n = 4, TBI n = 5. For imaging (J), sham n = 7, TBI n = 8. ● **Statistical Analysis:** Evaluated via one-way ANOVA with Tukey’s multiple-comparisons test for (A to D) and (H); and via two-way ANOVA with Šidák’s multiple-comparisons test in (F) and (J). ns = not significant; *p < 0.05; ***p < 0.001; ****p < 0.0001.

To isolate the cellular sources driving this chronic IFNγ cytokine pool, we paired flow cytometric profiling with *ex vivo* leukocyte stimulation (fig. S2D). Within the chronic thalamic niche (30 to 41 dpi), IFNγ-producing lymphocytes were markedly enriched compared to sham controls, whereas cells producing the type 2 (IL-5/13^+^) or type 3/17 (IL-17A^+^) effector cytokines were nearly undetectable (Fig. 2, E to G; fig. S2, E and F); a corresponding but more modest IFNγ-dominant lymphocyte profile was observed within the injured cortex (fig. S2, G and H). CD8^+^ T cells were the largest producers of IFNγ, constituting ∼80–90% of all IFNγ^+^ cells in the thalamus, with conventional CD4^+^ T cells accounting for the remaining fraction (Fig. 2, G and H, fig. S2, I to K).

We next mapped the spatial distribution of IFNγ–producing cells within the injured thalamus, using Tbet expression as a proxy for cells with the capacity to produce IFNγ (*31*). Confocal imaging of Tbet^zsGreen^ reporter mice (*35*) at 21 dpi revealed an accumulation of Tbet^zsGreen+^ cells compared to sham controls (Fig. 2I). Consistent with our regional T cell mapping (Fig. 1N), these Tbet-expressing cells clustered predominantly within the somatosensory thalamus, most abundantly in the VPL nucleus (Fig. 2J). By crossing this reporter system with a genetic CD8 lineage tracker (CD8^cre;tdTom+^), we confirmed that this spatial archetype consists of both conventional CD4^+^ T cells and lineage-tracked CD8^+^ T cells (Fig. 2K). Together, these data demonstrate that thalamic type 1 lymphocytes preferentially accumulate in the sensory thalamus several weeks after TBI and produce the effector cytokine IFNγ.

### Cortical TBI induces an interferon response within the ipsilateral sensory thalamus

Next, we used confocal microscopy to map the downstream microenvironmental consequences of this localized type 1 immune response. Consistent with our previous findings (*10*, *17*), cortical TBI increased the secondary glial reactivity across thalamic subregions, with accumulation of reactive astrocytes (GFAP^+^) and microglia/macrophages (Iba1^+^) at 14 dpi in the ipsilateral GABAergic reticular thalamic nucleus (nRT) and the glutamatergic VPL and ventroposteromedial (VPM) nuclei (fig. S3, A to C). The motor VAL nucleus remained devoid of reactive gliosis (fig. S3, B and C), supporting the hypothesis that thalamic inflammation occurs in the regions that are directly connected with the damaged cortex (*17*).

Because IFNγ initiates downstream transcription via canonical Janus kinase/Signal Transducer and Activator of Transcription (JAK/STAT) signaling (*36*), we examined the spatial expression of classical JAK/STAT-driven interferon-stimulated genes (ISGs). Major Histocompatibility Complex class II (MHC-II) and STAT1 proteins were upregulated after TBI; this upregulation was most pronounced in the injured VPL and lateral posterior/lateral dorsal (LP/LD) nuclei at 14 dpi (Fig. 3, A to C). To corroborate these observations, we utilized sensitive fluorescent reporter mice for two additional ISGs: Immunity-related GTPase family M member 1 (IRGM1^dsRed^) (*37*) and Interferon-Stimulated Gene 15 (ISG15^mGreenLantern(mGL)^) (*38*). Both IRGM1^dsRed+^ and ISG15^mGL+^ cells were elevated within sensory thalamic nuclei at 14 dpi (Fig. 3, D to G), demonstrating an active interferon response that maps precisely onto the spatial topography of the infiltrating type 1 lymphocytes (Fig. 2, I and J).

**Fig. 3.**
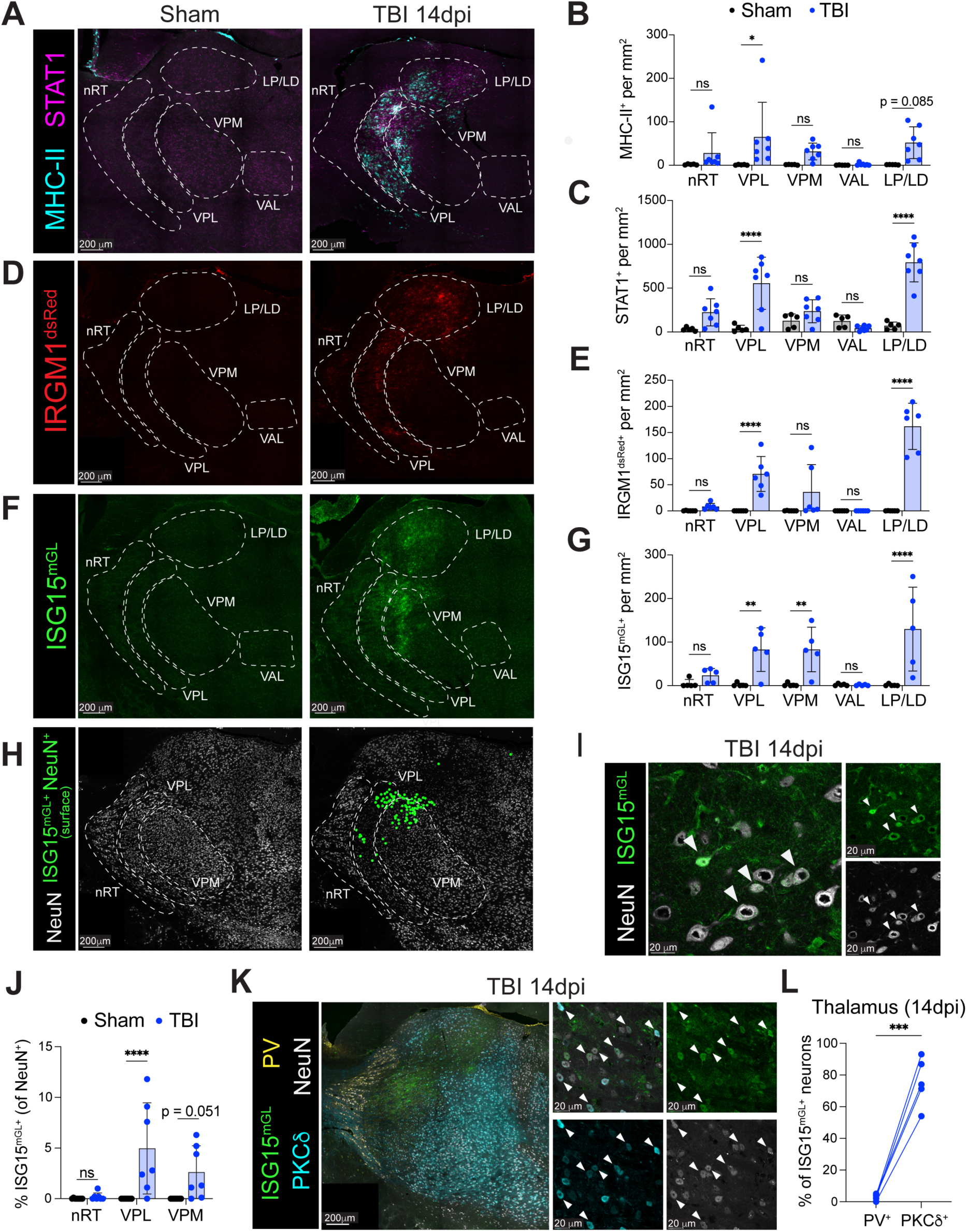
Cortical TBI induces a regional interferon response within the ipsilateral thalamus. **(A)** Representative confocal images of the ipsilateral thalamus from sham and TBI mice at 14 days post-injury (dpi), displaying immunofluorescence for MHC-II and STAT1. Anatomical subregions are delineated (dashed white lines) based on NeuN counterstaining (not shown). **(B and C)** Spatial quantification of MHC-II^+^ (B) and STAT1^+^ (C) cell density across thalamic subregions. **(D and E)** Representative confocal images (D) and corresponding regional quantification **(E)** of the thalamus from IRGM1 reporter mice (*Irgm1^dsRed^*) at 14 dpi, displaying IRGM1^dsRed+^ cell density. Anatomical subregions are outlined based on NeuN counterstaining (not shown). **(F and G)** Representative confocal micrographs (F) and corresponding regional quantification **(G)** of the thalamus from ISG15 reporter mice (*Isg15^mGL/+^*) at 14 dpi, displaying ISG15^mGL+^ cell density. **(H)** Confocal mapping of neuronal ISG15 expression, showing ISG15^mGL+^ NeuN^+^ neurons (green dots, 3D surface reconstruction) mapped over native NeuN staining across demarcated thalamic subregions. **(I)** High-magnification confocal images of the ipsilateral thalamus from an *Isg15^mGL/+^* reporter mouse (14 dpi) displaying merged (left) and individual (right) channels for ISG15^mGL^ and NeuN. White arrowheads indicate ISG15-expressing neurons. **(J)** Frequency of ISG15^mGL+^ neurons (calculated as a percentage of total NeuN^+^ cells) within designated thalamic subregions. **(K)** High-resolution confocal characterization of neuronal subtypes in the ipsilateral thalamus (14 dpi), displaying ISG15^mGL^, Parvalbumin (PV), Protein Kinase C delta (PKCδ), and NeuN. Insets (right) show high-magnification merged and single-channel views. White arrowheads mark ISG15^mGL+^ PKCδ^+^ NeuN^+^ excitatory neurons. **(L)** Proportion of ISG15^mGL+^ neurons that are either PV^+^ (inhibitory) or PKCδ^+^ (excitatory) across the ipsilateral thalamus at 14 dpi. Solid lines connect paired cellular frequencies from individual mice. **Data and Statistics:** Bar graphs and data points represent mean ± SD from individual mice (biological replicates). ● **Sample Sizes:** For MHC-II and STAT1 mapping (B and C), sham n = 5, TBI n = 7. For IRGM1^dsRed^ mapping (E), sham n = 6, TBI n = 6. For ISG15^mGL^ and neuronal mapping (G and J), sham n = 8, TBI n = 7. For neuronal subtype analysis (L), TBI n = 5. ● **Statistical Analysis:** Statistical significance was determined via two-way ANOVA with Šidák’s multiple comparisons test for all spatial mappings (B, C, E, G, J), and via a paired two-tailed Student’s *t*-test for within-subject subtype comparisons (L). ns = not significant; *p < 0.05; **p < 0.01; ***p < 0.001; ****p < 0.0001.

To determine the cellular architecture of this interferon response signature, we co-stained with cell type-specific markers. While MHC-II and STAT1 primarily colocalized with Iba1^+^ myeloid cells (fig. S3D), the reporters revealed broad multi-lineage responses: IRGM1^dsRed^ mapped predominantly onto GFAP^+^ astrocytes and OLIG2^+^ oligodendrocyte-lineage cells (fig. S3, E and F), whereas ISG15^mGL^ tracked with astrocytes and occasional microglia (fig. S3, G and H). Strikingly, ISG15^mGL^ expression was also observed within NeuN^+^ neurons within the injured somatosensory VPL nucleus, which was elevated over sham baselines (Fig. 3, H to J, fig. S3, G and H). Most (∼70%) interferon-responsive neurons across the injured thalamus were Protein Kinase C δ (PKCδ)^+^ excitatory neurons, while fewer than 10% were Parvalbumin (PV)^+^ inhibitory neurons (Fig. 3, K and L), based on expression of these lineage-specific markers (fig. S3I). This interferon response within neurons is concordant with a re-analysis of our previous single-nucleus RNA sequencing dataset (*10*), which confirmed that the heterodimeric receptor subunits *Ifngr1* and *Ifngr2* are expressed by thalamic excitatory neurons and microglia (fig. S3I). Thus, post-traumatic secondary injury drives a widespread parenchymal interferon response that includes thalamic neurons.

### Enhanced thalamic type 1 lymphocyte–IFNγ responses accompany seizure resilience after TBI

To determine the functional contribution of thalamic infiltrating lymphocytes and their IFNγ production on post-traumatic network instability, we first targeted CD4^+^ T cells (including T_regs_), depleting them subacutely following cortical injury using an antibody-mediated approach (Fig. 4, A and B) (*39*). CD4^+^ T cell depletion drove a localized, compensatory expansion of thalamic type 1 lymphocytes, including an increase in NK cells and a trending expansion of CD8^+^ T cells (Fig. 4, C to E). This lymphoid expansion was absent within the blood and perilesional cortex, despite systemic CD4^+^ T cell depletion (fig. S4, A to F), highlighting the anatomical specificity of this effect.

**Fig. 4.**
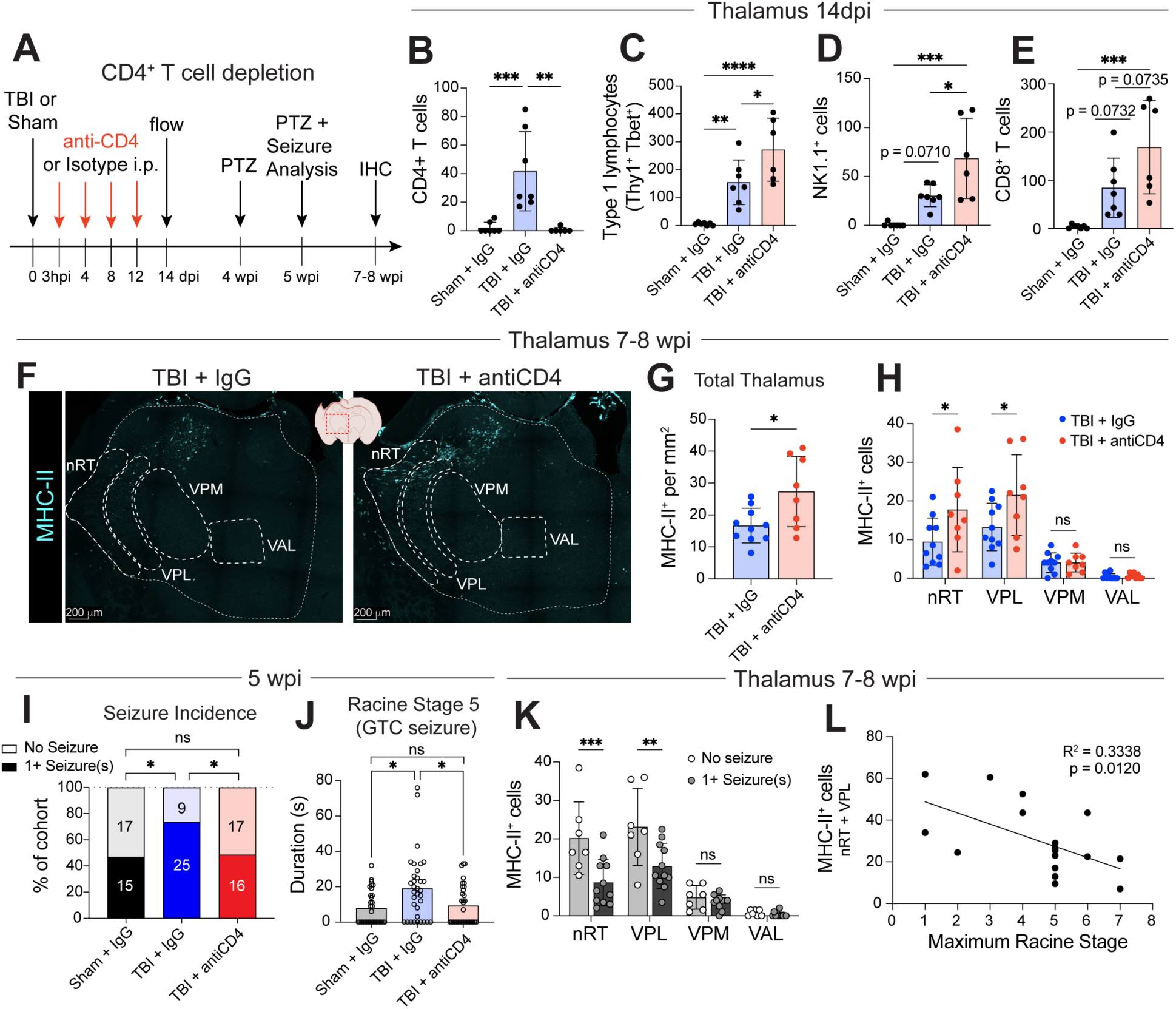
Depletion of CD4^+^ T cells dampens post-traumatic seizure susceptibility while amplifying thalamic IFNγ pathways. **(A)** Experimental workflow illustrating systemic anti-CD4 antibody (or IgG2b isotype) administration following sham or TBI surgeries and PTZ challenge paradigm to assess seizure susceptibility. **(B)** Number of thalamic CD4^+^ T cells, confirming CD4^+^ depletion with anti-CD4 treatment. **(C to E)** Number of infiltrating thalamic Thy1.2^+^ Tbet^+^ type 1 lymphocytes (C), NK cells (NK1.1^+^) (D), and CD8^+^ T cells (E). **(F)** Representative confocal images of the ipsilateral thalamus from IgG-or anti-CD4-treated TBI mice depicting MHC-II immunofluorescence. Anatomical subregions are outlined based on NeuN counterstaining (not shown) and the total ipsilateral thalamic region is delineated with a thin dotted line. **(G)** Density of MHC-II^+^ cells across the total ipsilateral thalamus of IgG- or anti-CD4-treated TBI mice. **(H)** Number of MHC-II^+^ cells across ipsilateral thalamic subregions from IgG- or anti-CD4-treated mice. **(I and J)** Seizure phenotype metrics showing the overall incidence of generalized tonic-clonic (GTC) seizures across experimental cohorts (I) and corresponding GTC seizure duration (Racine Stage 5) per animal (J). **(K)** Number of MHC-II^+^ cells across ipsilateral thalamus from mice stratified based on seizure presentation (0 vs. 1 or more GTC seizures. **(L)** Correlation between the maximum Racine stage reached per animal after PTZ challenge at 5 wpi and the sum of MHC-II^+^ cells within the ipsilateral nRT and VPL of each animal at 7-8 wpi. All mice received TBI. R^2^ = 0.3338, p = 0.0120. **Data and Statistics:** Data are mean ± SD unless otherwise indicated. Individual data points represent unique biological replicates (independent mice), except in (I and K) where individual data points represent single sequenced cells. ● **Sample size:** For (B to E), n = 7 per group. For (G and H), TBI+IgG n = 10, TBI+anti-CD4 n = 8. For (I and J), sham+IgG n = 32, TBI+IgG n = 34, TBI+anti-CD4 n = 33. For (K), no seizure n = 7, 1+ seizure(s) n = 11. For (L), n = 18. ● **Statistical Analysis:** Evaluated via one-way ANOVA with Tukey’s post-hoc test for (B to E), unpaired Student’s *t*-test for (G), two-way ANOVA with Šidák’s multiple comparisons test for (H and K), Fisher’s exact test for (I), Kruskall-Wallis test with Dunn’s multiple comparisons test for (J), and simple linear regression for (R). Statistics: ns = not significant, *p < 0.05, **p < 0.01, ***p < 0.001, ****p < 0.0001.

To determine if this thalamic type 1 lymphocyte expansion amplified downstream IFNγ signaling, we first established an *in vivo* transcriptomic fingerprint of IFNγ responsiveness in microglia and mapped this signature onto single-cell RNA sequencing (scRNA-seq) profiles of thalamic myeloid cells (CD11b^+^) at 14 dpi (fig. S4, G to L). scRNA-seq clustering resolved microglial activation states, including disease-associated (DAM) (*40*) and interferon-responsive (IRM) (*41*) clusters, that were expanded after TBI; notably, anti-CD4 treatment further enriched the frequency of DAM and IRM clusters (fig. S4, G to J) and the IFNγ signature of mature DAMs (fig. S4, K to M). We validated that anti-CD4 treatment increased the frequency of DAMs (CD11c^+^ CLEC7A^+^ microglia) via flow cytometry at 14 dpi (fig. S4N) and the abundance of thalamic IFNγ-stimulated MHC-II^+^ cells via microscopy at 7-8 wpi (Fig. 4, F to H). These data illustrate that anti-CD4 treatment enhances the expansion of total type 1 lymphocytes and the subsequent IFNγ response within the thalamus after TBI.

To test the functional impact of this heightened thalamic type 1 lymphocyte – IFNγ-response, mice were subjected to a chemoconvulsant challenge with the GABA_A_ receptor antagonist pentylenetetrazol (PTZ) to evaluate neuronal network excitability at 5 weeks post-injury (wpi) (*11*, *42*); seizure behavior was quantified and scored on a modified Racine scale (Fig. 4A, Table S1) (*42*, *43*). Elevations of thalamic type 1 lymphocytes and IFNγ signaling (driven by CD4^+^ T cell depletion) protected injured mice against post-traumatic network instability, reducing both the incidence and duration of generalized tonic-clonic (GTC) seizures compared to isotype-treated injured controls (Fig. 4, I and J, fig. S4, O to Q). This heightened thalamic interferon state directly correlated with functional circuit resilience: post-TBI mice that remained entirely resilient to generalized seizures had more MHC-II^+^ cells within the somatosensory thalamic nuclei as compared to their seizure-susceptible counterparts (Fig. 4K). Moreover, the number of MHC-II^+^ cells within the nRT and VPL of each mouse post-TBI was inversely correlated with the maximum Racine stage each mouse attained after PTZ challenge (Fig. 4L), revealing that heightened IFNγ responses within the somatosensory thalamus are associated with less severe seizure behaviors. Together, these findings reveal a competitive neuroimmune axis wherein CD4^+^ T cells counter-regulate a protective, type 1 lymphocyte-IFNγ cascade that restricts network hyperexcitability after TBI.

### IFNγ signaling to neurons suppresses post-traumatic hyperexcitability

To determine whether endogenous IFNγ signaling directly constrains post-traumatic network instability, we subjected global IFNγ receptor 1 knockout mice (*Ifngr1^−/−^*) (*44*) and wild-type controls to TBI with chemoconvulsant challenge at 5 wpi (Fig. 5A). Global loss of IFNγ signaling exacerbated seizure susceptibility: *Ifngr1^−/−^* mice exhibited a marked increase in the incidence and duration of GTC seizures, the overall severity of seizure behavior, and the seizure-induced mortality rate compared to wild-type injured counterparts (Fig. 5, B to D, fig. S5, A to C). These data demonstrate that IFNγ signaling acts as an indispensable endogenous brake against post-traumatic seizure risks.

**Fig. 5.**
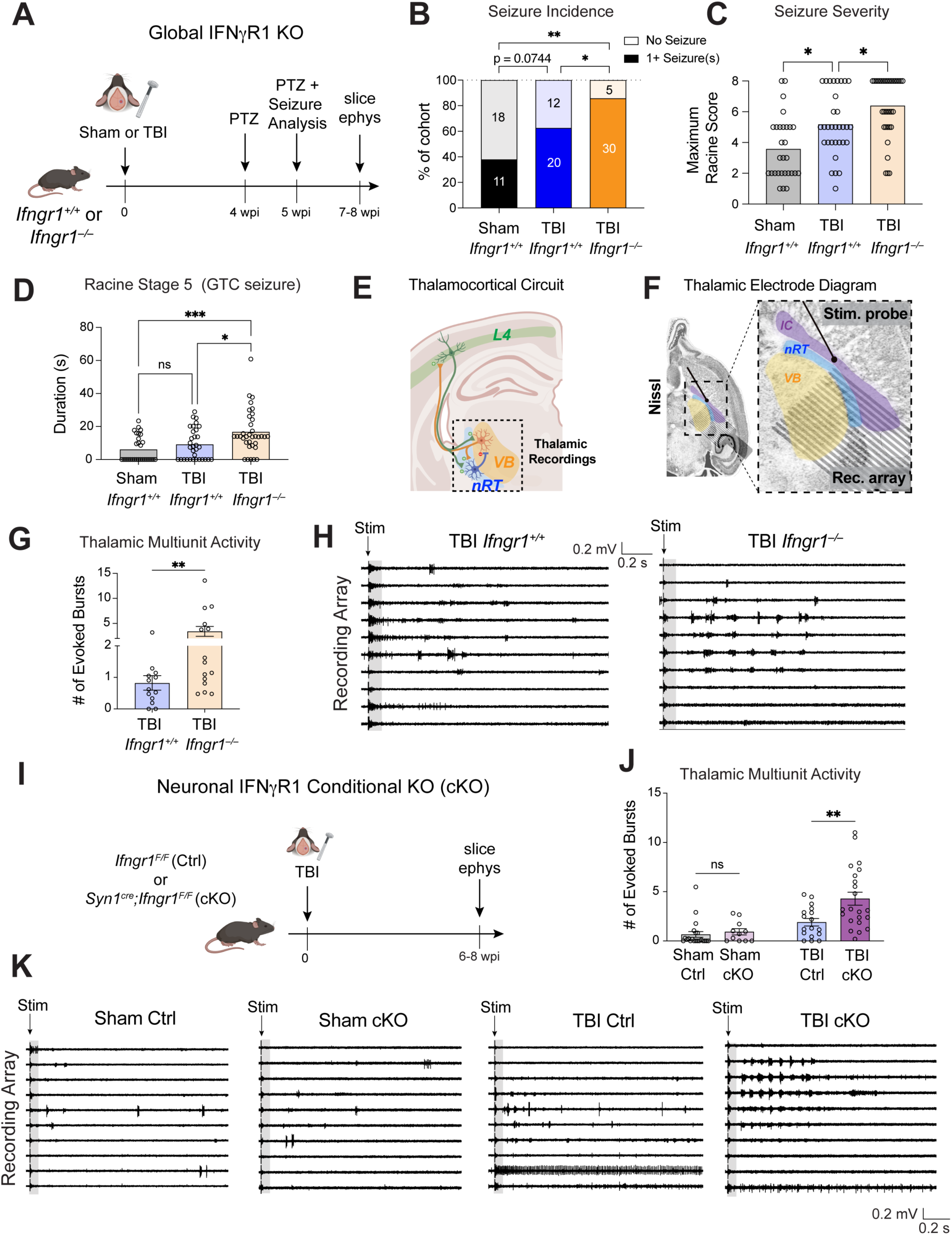
Endogenous IFNγ signaling in neurons is required to suppress post-traumatic hyperexcitability. **(A)** Experimental paradigm evaluating seizure susceptibility and network hyperexcitability in global IFNγR1 knockout (*Ifngr1^-/-^*) mice or wild-type (*Ifngr1^+/+^*) controls. **(B to D)** Seizure phenotype metrics evaluated at 5 weeks post-injury (wpi), showing the percentage of animals experiencing one or more generalized tonic-clonic (GTC) seizures (B), the corresponding maximum Racine severity scores reached per animal (C), and corresponding GTC seizure duration (Racine Stage 5) per animal (D). **(E)** Diagram of thalamocortical circuit highlighting relevant circuitry for thalamic recordings. **(F)** Schematic of recording and stimulating electrodes’ placements in the thalamus and internal capsule, respectively. **(G and H)** Evoked network activity in the thalamus displaying absolute numbers of evoked thalamic bursts (G) and representative thalamic recordings across the indicated experimental groups (H). **(I)** Workflow to evaluate functional network hyperexcitability following conditional, neuron-specific deletion of the IFNã receptor (Syn1^ΔIfngr1^*; (Syn1^cre^; Ifngr1^F/F^*)) compared against Cre-negative littermate controls (*Ifngr1^F/F^*) after TBI. **(J and K)** Evoked network activity in conditional neuronal knockouts (cKOs) and controls (Ctrls) displaying absolute numbers of evoked thalamic bursts (J) and representative thalamic recordings across the indicated experimental groups (K). **Data and Statistics:** Data are means in (C and D) and mean ± SEM in (G and J). Data points represent individual mice in (C and D) and brain slices in (G and J). IC = internal capsule. ● **Sample size:** For (B to D), sham *Ifngr1^+/+^* n = 29; TBI *Ifngr1^+/+^* n = 32; TBI *Ifngr1^-/-^* n = 35. For (G), TBI WT n = 13 slices from n = 8 mice; TBI KO n = 15 slices from n = 10 mice. For (J), Sham Ctrl n = 21 slices from n = 8 mice; Sham cKO n = 11 slices from n = 6 mice; TBI Ctrl n = 17 slices from n = 8 mice; TBI cKO n = 22 slices from n = 11 mice. ● **Statistical Analysis:** Evaluated via Fisher’s exact test for (B), Kruskall-Wallis test with Dunn’s multiple comparisons test for (C and D), Mann-Whitney U-test for (G), and two-way ANOVA with Šidák’s multiple comparisons test (J). Statistics: ns = not significant, *p < 0.05, **p < 0.01.

We next utilized *ex vivo* slice electrophysiology to dissect the circuit-level mechanisms underlying this protection, recording across the reciprocal loop linking the somatosensory ventrobasal (VB) thalamus and layer 4 of the somatosensory cortex (Fig. 5E); this sleep-spindle-generating circuit is of particular interest because it generates epileptiform discharges (*10*), a key aspect of network dysfunction that is predictive of long-term epilepsy (*45*, *46*). Multiunit activity recordings within the VB thalamic complex following afferent stimulation of the internal capsule (Fig. 5F) revealed a profound increase in evoked burst discharges in post-TBI *Ifngr1^−/−^* mice compared to injured wild-type controls (Fig. 5, G to H). Parallel local field potential (LFP) recordings and current source density (CSD) analyses within the somatosensory cortex revealed a concurrent trend toward hyperexcitability that localized predominantly to thalamorecipient layer 4 (fig. S5, D to M). Endogenous IFNγ signaling is therefore required to dampen pathological hypersynchrony across the core loops of the thalamocortical circuit.

Given that thalamic neurons express the receptor complex and respond to IFNγ after TBI *in vivo* (Fig. 3, H to L; fig. S3, G to I), we hypothesized that IFNγ mediates these protective effects via direct neuronal engagement. To test this, we generated mice with conditional deletion of IFNγR1 specifically within neurons (*Syn1^cre^;Ifngr1^F/F^*, hereafter referred to as Syn1^ΔIfngr1^ conditional knockout (cKO) mice) (Fig. 5I) (*47*, *48*). At 5 wpi, functional circuit mapping using thalamic multi-unit recordings revealed that injured Syn1^ΔIfngr1^ cKO mice exhibited a pronounced increase in evoked burst discharges compared to Cre-negative littermate controls (Fig. 5, J and K). Collectively, these findings demonstrate that neuronal sensing of IFNγ suppresses pathological circuit hyperexcitability and protects the brain from post-traumatic epileptiform networks.

### IFNγ treatment reduces thalamocortical hyperexcitability, seizure risk, and mortality after TBI

Having established that neuronal IFNγ responses promote network stability, we reasoned that exogenous administration of the cytokine could therapeutically rescue post-traumatic circuit dysfunction. We first evaluated the acute network effects of recombinant IFNγ delivery using *ex vivo* slice electrophysiology 24 hours following a single systemic injection (Fig. 6A). Within the somatosensory thalamus, the pathological elevation of evoked burst discharges observed in injured, vehicle-treated controls was reversed by IFNγ treatment (Fig. 6, B and C). Complementary CSD profiling within the perilesional somatosensory cortex revealed a concurrent reduction in the amplitude of both presynaptic and postsynaptic current sources and sinks (fig. S6, A to E), demonstrating that systemic IFNγ delivery is sufficient to reset post-traumatic thalamocortical hyperexcitability.

**Fig. 6.**
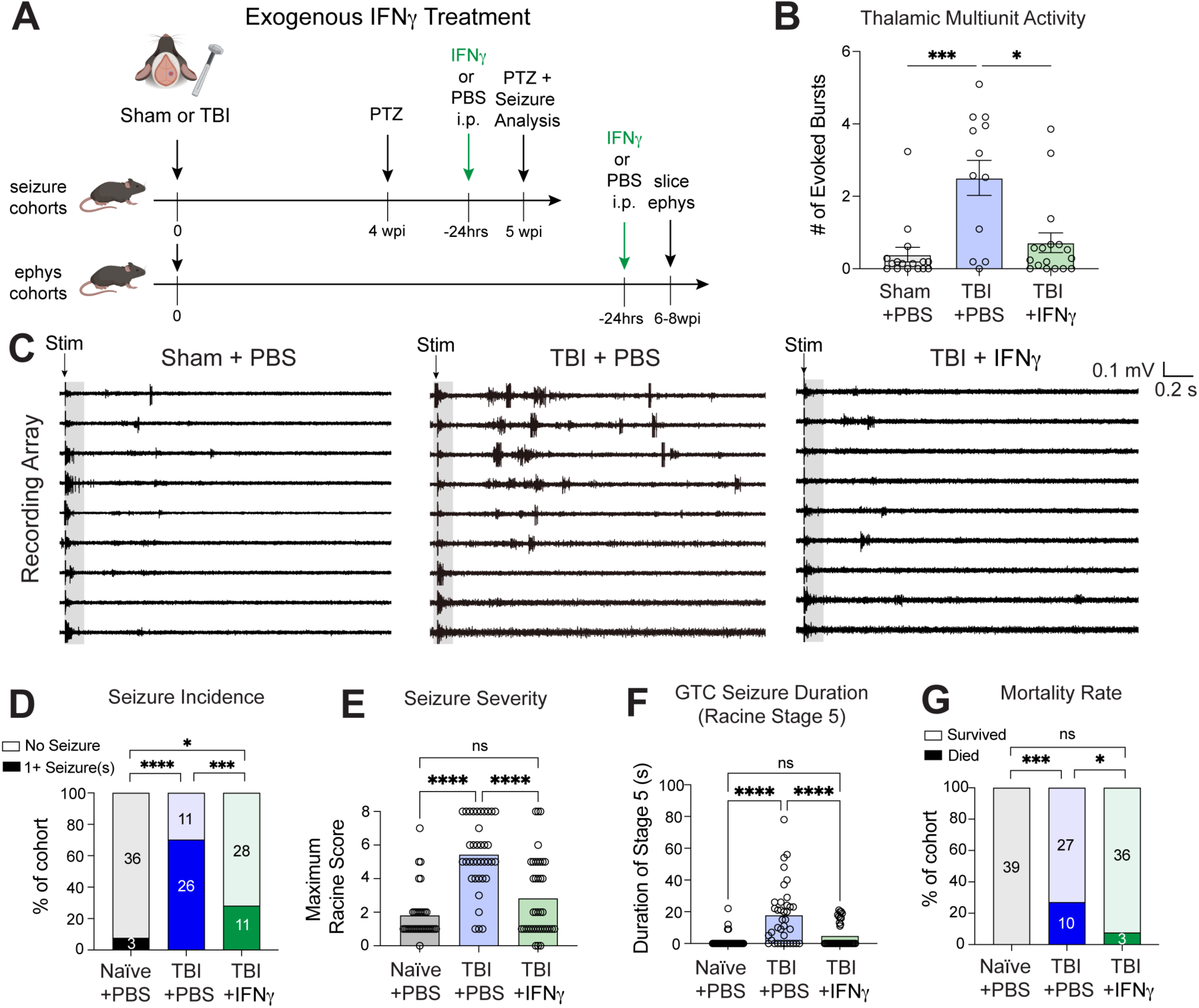
Exogenous IFNγ administration mitigates post-TBI thalamocortical hyperexcitability and protects against seizure risks and mortality. **(A)** Experimental paradigms for assessing long-term seizure susceptibility (top) and functional network excitability (bottom) in mice following TBI and IFNγ (or PBS vehicle) administration *in vivo* (24 hours before assessment). **(B and C)** Evoked network activity following TBI and IFNγ (or PBS vehicle) administration displaying absolute numbers of evoked thalamic bursts (B) and representative thalamic recordings across the indicated experimental groups (C). **(D)** Percentage of mice that experienced one or more generalized tonic-clonic (GTC) seizures following PTZ challenge. **(E and F)** Seizure phenotype metrics after PTZ challenge, showing the total duration of GTC seizures (Racine Stage 5) per mouse (E) and corresponding maximum convulsive severity scores reached per mouse (F). **(G)** Mortality rates calculated as the percentage of mice that died within 20 minutes of PTZ challenge. **Data and Statistics:** Data are mean ± SEM in (B) and means in (E and F). Data points represent individual brain slices in (B) and individual mice in (E and F). ● **Sample size:** For (B), Sham+PBS n = 16 slices from n = 8 mice; TBI+PBS n = 13 slices from n = 8 mice; TBI+IFNγ n = 17 slices from n = 8 mice. For (D to G), mice per condition (pooled from 2 independent cohorts): naïve + PBS n = 39, TBI + PBS n = 37, TBI + IFNγ n = 39. ● **Statistical Analysis:** Evaluated via Kruskall-Wallis test with Dunn’s multiple comparisons test for (B, E, and F), and Fisher’s exact test for (D and G). Statistics: ns = not significant, *p < 0.05, ***p < 0.001, ****p < 0.0001.

To determine if this electrophysiological normalization translates into seizure protection, we administered a single dose of IFNγ or vehicle control to chronically injured mice 24 hours prior to chemoconvulsant challenge at 5 wpi (Fig. 6A). While vehicle-treated injured mice exhibited a high incidence of GTC seizures, acute IFNγ treatment profoundly blunted this post-traumatic seizure risk (Fig. 6D). Beyond reducing absolute seizure incidence, IFNγ therapy curtailed GTC seizure duration and overall paroxysmal severity, shifting Racine scores toward milder phenotypes and restricting time spent in advanced convulsive stages (Fig. 6, E and F, fig. S6, F to M). Most notably, a single treatment with IFNγ protected mice against post-traumatic mortality following PTZ challenge, reducing the death rate from 27.0% in vehicle controls to 7.7% (Fig. 6G). Thus, acute therapeutic engagement of the type 1 IFNγ axis is sufficient to reverse established network hyperexcitability, providing robust protection against post-traumatic epileptiform networks and related mortality.

## Discussion

Our findings establish that TBI prompts a delayed, anatomically-targeted infiltration of peripheral lymphocytes that actively preserves circuit stability within the chronic post-injury landscape. Rather than driving secondary immunopathology, these tissue-resident type 1 lymphocytes – comprised of CD8^+^ T cells, CD4^+^ T cells, and NK cells – establish a chronic protective interferon-γ (IFNγ) microenvironment across the damaged thalamocortical system. Selective depletion of CD4^+^ T cells uncouples an endogenous immune checkpoint, de-repressing this type 1 cascade to augment IFNγ responses and shield the brain from network failure. We demonstrate that IFNγ signaling is an endogenous brake on thalamocortical network instability that operates through direct, cell-autonomous IFNγ signaling to neurons. Crucially, we determine that a single systemic dose of recombinant IFNγ can acutely reverse established hypersynchrony and reduce post-traumatic seizure risks and seizure-related mortality. Together, these data reveal that the adaptive immune response installs a potent homeostatic brake on neural circuit hyperexcitability that can be therapeutically leveraged long after the primary insult.

The mechanisms governing the recruitment, antigen-specificity, and spatial organization of central nervous system (CNS)-resident lymphocytes after mechanical trauma remain a major frontier in neuroimmunology. While initiation of a type 1 IFNγ-dominant immune response appears conserved across multiple models of CNS injury (*27*, *28*), the cellular and molecular factors directing this polarization and subsequent recruitment of peripheral lymphocytes to the CNS remain unknown. Following TBI, central T cell seeding often requires peripheral lymph node priming against shed CNS autoantigens (*18*, *27*); whether the thalamic T cell repertoire after TBI recognizes a distinct, injury-induced antigenic landscape remains to be determined.

While neurotrauma causes broad blood-brain barrier disruption, the highly regionalized accumulation of type 1 lymphocytes within the sensory thalamus suggests an orchestrated homing process to damaged areas. In other models of neurological disease, microglial-derived chemokines direct CD8^+^ T cell homing (*26*, *49–51*) and fibroblast-derived chemokines recruit T cells to the CNS (*28*, *52–54*). Our transcriptomic data suggests that microglia after TBI express several putative T cell recruiting chemokines, raising the possibility that reactive microglia are responsible for the regional recruitment of T cells to the damaged sensory thalamus. Crucially, this regional lymphoid distribution provides a mechanistic rationale for the subregional vulnerabilities characteristic of the secondarily injured thalamus. We previously demonstrated that chronic post-traumatic dysfunction is explicitly manifested by a loss of GABA_A_-mediated synaptic inhibition within the reticular thalamic nucleus (nRT) (*10*, *13*, *17*). Because the protective IFNγ cascade identified here is heavily skewed toward the neighboring glutamatergic ventroposterolateral (VPL) nucleus, a localized deficit in lymphoid-derived paracrine factors within the nRT may directly underlie its profound chronic vulnerability.

While IFNγ has been implicated as a behavioral modulator (*55*, *56*), our data demonstrate a clear functional requirement of IFNγ neuromodulation in stabilizing the post-traumatic thalamocortical circuit. This finding aligns with evidence that neurons across species can respond to IFNγ and upregulate canonical interferon-stimulated genes (ISGs) (*57–61*). Mechanistically, IFNγ signaling bolsters synaptic inhibition in healthy circuits by driving the surface expression and phosphorylation of GABA_A_ receptors (*62–64*). In contrast, the classical complement pathway exacerbates post-traumatic pathology within the same circuit: we showed that microglial-derived complement factor C1q accumulates chronically within the post-injury thalamocortical network and promotes GABAergic nRT neurodegeneration, sleep architecture disruption, and the emergence of focal epileptiform spikes (*10*). Our finding that IFNγ signaling instead acts as an indispensable homeostatic brake highlights a nuanced duality in the post-traumatic neuroimmune landscape: while chronic, microglial-driven innate complement cascades can be profoundly maladaptive, a concurrent or subsequent wave of adaptive, lymphoid-derived type 1 cytokines is endogenously deployed to constrain runaway network synchrony. Whether this protective braking mechanism relies on canonical JAK/STAT-mediated transcription or operates via non-transcriptional, localized post-translational regulation of synaptic machinery (*59*, *65–67*) remains an essential question for future dissection.

Beyond direct neuronal network stabilization, IFNγ concurrently shapes the activation states of non-neuronal parenchymal cells, particularly microglial states. Interferon-responsive (IRM) and disease-associated (DAM) microglia exhibit highly augmented phagocytic capacities, which can alter the disease course by accelerating the clearance of cellular debris, stressed neurons, or refining aberrant synaptic structures (*40*, *41*, *68–72*). Indeed, elevated DAM abundance correlates with attenuated seizure severity in alternative models of epilepsy (*71*). It is also probable that a cooperative neuron-glia dialogue is coordinated by this cytokine cascade; a CD8-IFNγ-neuron-microglia axis has been shown to mediate synaptic and behavioral alterations during neuroinflammation (*72*). Dissecting the precise contributions between direct neuronal plasticity and microglial-mediated circuit remodeling will further clarify how this neuroimmune axis coordinates chronic network resilience.

Translational strategies for traumatic brain injury and post-traumatic epilepsy have historically focused on broadly suppressing neuroinflammatory responses to limit secondary injury. Consistent with this approach, IFNγ has often been viewed as a pathogenic inflammatory mediator based on its elevation in epileptic tissue, association with disease severity in human epilepsy, and evidence implicating interferon signaling in seizure-promoting inflammatory cascades (*73–79*). Although recent work demonstrated that exogenous IFNγ can reduce induced seizures (*56*, *80*), its endogenous role within injured neural circuits has remained poorly understood. Here, we identify a previously unrecognized protective function for endogenous IFNγ after traumatic brain injury. Rather than amplifying pathology, thalamic IFNγ signaling limits post-traumatic thalamocortical hyperexcitability and hypersynchrony, reduces seizure susceptibility, and protects against seizure-associated mortality. These findings reveal that the injured brain actively recruits type 1 lymphocytes to establish a specialized neuroimmune niche in which IFNγ acts on surviving neurons to promote long-term circuit stability and resilience. More broadly, our results challenge the prevailing assumption that post-traumatic immune responses are detrimental and suggest that spatially restricted immune signaling can serve an adaptive role in maintaining neural function after injury.

## Acknowledgements

We are grateful to the A.V. Molofsky, A.B. Molofsky, and J.T. Paz labs for helpful comments on the manuscript. We thank Dr. J. Nilsson and V. DeNittis for assistance with mouse perfusions and tissue preparation. We thank Dr. Jinfang Zhu (NIAID) for generous sharing of Tbet^zsGreen^ mice and Dr. Michael Rosenblum (UCSF) for generous sharing of CD8^cre;tdTom^ mice. We thank Dr. Sarah Anderson for advice on immunostaining brain tissue from ISG15^mGL^ mice. We thank Dr. Matthew Spitzer (UCSF) for generous sharing of IRGM1^dsRed^ mice that were obtained from the Mutant Mouse Resource and Research Center (MMRRC). We thank the UCSF Parnassus Flow Core (RRID:SCR_018206 supported in part by Grant NIH P30 DK063720 and by the NIH S10 1S10OD021822-01), the UCSF Biological Imaging and Development Colab (BIDC), the Weill Innovation Core (WIC) Advanced Light Microscopy Core at the Weill Institute for Neurosciences, the UCSF Center for Advanced Technology (CAT), and the UCSF and Gladstone Institutes Laboratory Animal Resource Center (LARC) for instruments, services, and support. Schematics were created in part using https://BioRender.com.

## Author Contributions

Conceptualization: NMM, AC, AVM, ABM, JTP

Methodology: NMM, AC, NAE-C, LCD, RTH, AVM, ABM, JTP

Investigation: NMM, AC, NAE-C, LCD, RTH, JJB, NJS, AM

Visualization: NMM, AC, LCD, AVM, ABM, JTP

Funding acquisition: AVM, ABM, JTP Supervision: AVM, ABM, JTP

Writing – original draft: NMM, AC, AVM, ABM, JTP

Writing – review and editing: all authors

## Data, code, and materials availability

Tbet^zsGreen^ mice were obtained via a material transfer agreement and IRGM1^dsRed^ mice were obtained via MMRRC conditions of use agreement. Transcriptional datasets will be available via GEO (accession number to be provided upon acceptance of manuscript).

## MAIN TEXT METHODS

### Study design and animals

All animal procedures were conducted in accordance with the National Institutes of Health (NIH) guidelines for the care and use of laboratory animals and were approved by the Institutional Animal Care and Use Committees (IACUCs) at the University of California, San Francisco and Gladstone Institutes. Precautions were taken to minimize stress and the number of animals used in each set of experiments. Male and female C57BL/6J mice (8-12 weeks old) were utilized. Various transgenic reporter and conditional knockout lines (including *Tbx21^zsGreen^, Cd8a^E8i−cre^, Rosa26^LSL−tdTom^, Isg15^mGL^, Irgm1^dsRed^, Ifngr1^−/−^, Ifngr1^Flox^,* and *Syn1^cre^*) were bred in-house to map specific lymphocyte and interferon-responsive populations. Additional details are described in supplementary materials and methods.

### Controlled Cortical Impact

Traumatic brain injury was induced via controlled cortical impact (CCI) (*10*). Briefly, adult mice were anesthetized with 2–5% isoflurane and a 3-mm craniotomy was centered over the right primary somatosensory (S1) cortex centered at-1 mm posterior from bregma, +3 mm lateral from the midline. A stereotaxic electromagnetic impactor (Impact One Stereotaxic Impactor for CCI, Leica Microsystems) delivered a contusion at a 0.8 mm depth from dura, 3 m/s velocity, and 300 ms dwell time. Sham mice received identical anesthesia and a trace craniotomy without cortical impact. Additional details are described in supplementary materials and methods.

### In vivo pharmacological interventions

For CD4^+^ T cell depletion, mice received an intraperitoneal (i.p.) injection of anti-mouse CD4 antibody (clone GK1.5) (*39*) or an isotype control (IgG2b, clone LTF-2) 3 hours post-injury, followed by maintenance booster doses every 4 days for two weeks. For cytokine rescue experiments, mice received a single i.p. injection of recombinant mouse IFNγ (10 μg) (*81*) 24 hours prior to proconvulsant challenge or electrophysiological recording. Additional details are described in supplementary materials and methods.

### Proconvulsant challenge and behavioral analysis

Seizure susceptibility was assessed using the chemoconvulsant pentylenetetrazol (PTZ) as previously described (*11*, *42*). At 4 weeks post-injury, mice received a sub-convulsive priming dose of PTZ (45 mg/kg, i.p.). One week later, mice were challenged with a second 45 mg/kg PTZ dose and monitored via continuous video recording for 20 minutes. Seizure severity, incidence of generalized tonic-clonic (GTC) seizures, and mortality were quantified offline using a modified Racine scale (*42*, *43*) (see Table S1) by blinded investigators. Additional details are described in supplementary materials and methods.

### Tissue processing, flow cytometry, and imaging

For flow cytometry, mechanically dissociated brain tissues (cortex, hippocampus, thalamus) were subjected to Percoll density gradient centrifugation to isolate mononuclear cells as previously described (*82*). Cells were stained for surface and intracellular markers (including transcription factors and intracellular cytokines following *ex vivo* PMA/ionomycin stimulation) and analyzed on a BD Fortessa cytometer. For immunofluorescence, PFA-fixed brains were cryosectioned, stained with primary and secondary antibodies, and imaged using high-resolution laser scanning confocal microscopy (Nikon A1R, Leica Stellaris 8, or Zeiss LSM800). Cellular quantification and 3D reconstructions were performed using Imaris software. Thalamic cytokine levels were quantified from homogenized micro-dissected tissues using a Luminex multiplex array. Additional details are described in supplementary materials and methods.

### Single-cell RNA sequencing

Myeloid cells (CD11b^+^) were isolated via MACS bead enrichment. For single-cell RNA sequencing (scRNAseq), libraries were generated using the 10x Genomics Chromium platform and sequenced on an Illumina NovaSeq. Data were processed using Cell Ranger and analyzed via Seurat, including calculation of IFNγ signature scores. For bulk RNA sequencing, microglia were sorted from juvenile mice 22 hours after an intracerebroventricular (i.c.v.) injection of recombinant IFNγ, and libraries were processed using the Lexogen QuantSeq kit and analyzed via DESeq2. Additional details are described in supplementary materials and methods.

### Electrophysiology

#### Ex vivo thalamic electrophysiology

To evaluate intra-thalamic microcircuit excitability, horizontal brain slices (400 μm) preserving the connectivity between the reticular thalamic nucleus (nRT) and the somatosensory ventrobasal thalamus (containing VPL and VPM) were prepared from transcardially perfused mice (*11*). Extracellular multi-unit activity (MUA) was recorded across the nRT and VB using a 16-channel linear multi-electrode array (Neuronexus) in an interface chamber. Evoked circuit responses were elicited via bipolar electrical stimulation of the internal capsule (*11*).

#### Ex vivo neocortical laminar electrophysiology

To map functional synaptic alterations across cortical laminae, extracellular local field potentials (LFPs) were recorded in the perilesional cortex using a 16-channel linear electrode array spanning layers 1 through 6. Synaptic responses were evoked via bipolar electrical stimulation of the underlying white matter (*83*). To dissect the distinct biophysical components of the network response, pharmacological bath application of glutamatergic antagonists and sodium channel blockers was used to isolate pre-versus post-synaptic potentials (*83*). The spatiotemporal location, direction, and magnitude of transmembrane synaptic currents were mathematically resolved by calculating the second spatial derivative of the laminar LFPs to generate Current Source Density (CSD) profiles (*83*, *84*) (see further details in statistical analysis below and the supplementary materials and methods).

### Statistical Analysis and Software Platforms

Extracellular wave data and multi-unit spiking metrics were visualized and analyzed using Spike2 (v9.0, Cambridge Electronic Design; RRID:SCR_000903). Analysis of flow cytometric data was performed using FlowJo (v10, BD). Confocal image processing, surface area calculations, and volumetric 3D reconstructions were performed using Bitplane Imaris (v9.5.1, Andor Technology). Custom data restructuring, matrix manipulation, and two-dimensional Current Source Density (CSD) spatial derivative plotting were executed via MATLAB (vR2023b, MathWorks; RRID:SCR_001622; CSD code kindly provided by the Huguenard lab) (*84*). Statistical analyses were conducted in SigmaPlot (v15.0, Systat Software; RRID:SCR_003210) and OriginPro (v2024, OriginLab; RRID:SCR_014212). All downstream bioinformatic tests, histological group quantifications, and survival metrics were analyzed using GraphPad Prism (v11.0.2, GraphPad Software; RRID:SCR_002798). Data are presented as mean ± SD, unless stated otherwise and significance criteria were evaluated at α=0.05. Parametric and non-parametric tests were chosen as appropriate and were reported in figure legends.

