## Supplementary Files for "Type 1 lymphocytes accumulate in the thalamus and produce interferon-gamma to restrict seizure susceptibility after brain injury"

Figs. S1 to S6

Tables S1-S4

References (81-98)

### **Materials and Methods**

**Animals**

Adult C57BL/6J mice (8–12 week old) (JAX #000664) were purchased from JAX and used for most experiments. Mice weighed 22–28 g at the time of TBI/sham surgery. Both female and male mice were used in scRNA-seq and reporter imaging experiments whereas all other experiments used only male mice.

*Tbx21^zsGreen^* (Tbet^zsGreen^) (MGI #5690118) (*35*) were generously shared by Dr. Jinfang Zhu (NIAID). *Cd8a^E8i-cre^* (JAX #008766) (*85*) and *Rosa26^LSL-tdTom^* (Ai14, JAX #007914) (*86*) mice were received from Dr. Michael Rosenblum (UCSF) and intercrossed in-house for imaging experiments. *Isg15^mGL^* reporter mice (IFN-brite, JAX #041458)(*38*) were generated and bred in-house. *Irgm1^dsRed^* reporter mice (M1Red, #06913-JAX) (*37*) were obtained from the Mutant Mouse Resource and Research Centers (MMRRC) (submitted by Dr. Alan Sherr and Dr. Lionel Feigenbaum) via collaboration with Dr. Matthew Spitzer (UCSF). IFNγR1 knockout mice (*Ifngr1^–/–^*, JAX #003288) (*44*), IFNγR1^Flox^ mice (*Ifngr1^Flox^*, JAX #025394) (*47*), and Syn1^cre^ mice (JAX #03966) (*48*) were purchased from JAX, intercrossed, and bred in-house for experiments.

**Controlled cortical impact (CCI)**

Controlled cortical impact (CCI) was induced in the somatosensory cortex as previously described (*10*). Briefly, in isoflurane-anesthetized adult mice, a 3-mm diameter craniotomy was performed over the right somatosensory (S1) cortex (centered at -1 mm posterior to Bregma and 3 mm lateral to the midline). TBI was performed with a CCI device (Impact One Stereotaxic Impactor for CCI, Leica Microsystems) equipped with a metal piston using the following parameters: 3 mm tip diameter, 18° angle, 0.8 mm depth from the dura, 3 m/s velocity, and 300 ms dwell time. Sham mice received identical anesthesia and scalp incision but underwent a sham 3-mm craniotomy in which the drill bit was used to trace the craniotomy without creating a bone flap; no cortical impact was delivered. Mice received buprenorphine (0.05 mg/kg) at the end of the surgery in accordance with Institutional Animal Care and Use Committee (IACUC)-approved protocols.

**PTZ challenge and seizure analysis**

To assess seizure susceptibility after TBI in a high-throughput manner, we evaluated behavioral seizures induced by intraperitoneal (i.p.) administration of the convulsant pentylenetetrazol (PTZ; Tocris; 45 mg/kg). The PTZ dose was selected based on prior studies for C57BL/6 mice in our group and others.(*11*, *42*) Four weeks after TBI/sham surgery mice received a first (“priming”) dose of PTZ (45 mg/kg) without video recording. One week later, mice received a second dose of PTZ (45 mg/kg) and were video-recorded for 20 minutes after injection. Videos were analyzed offline for the incidence of generalized tonic-clonic (GTC) seizures, mortality, and seizure severity using the modified Racine scale (*42*, *43*) (Table S1). All mice used for seizure analysis were male.

**In vivo antibody-mediated depletion**

Rat anti-mouse CD4 antibodies (250 μg, clone GK1.5, BioXCell) or the corresponding isotype control rat IgG2b antibodies (250 μg, clone LTF-2, BioXCell) were diluted in sterile PBS buffer (BioXcell) and administered intraperitoneally 3 hours after TBI or sham surgery. Both antibody-treated cohort and isotype control-treated mice received booster injections every 4 days (250 μg in sterile PBS) for two constitutive weeks to maintain CD4 T cell depletion.

**In vivo cytokine treatment**

Recombinant mouse IFNγ (carrier-free, BioLegend) was reconstituted in sterile double-distilled water and freshly diluted in 1X Dulbecco’s phosphate-buffered saline (DPBS) before administration. IFNγ (10 μg per animal in a total volume of 200 μL) was administered i.p. 24 hours before PTZ challenge recording or harvest for *ex vivo* electrophysiology. Vehicle-treated TBI and sham mice received 200 μL of 1X DPBS.

**Electrophysiology**

***Brain slice preparation***

Slice preparation was performed using neuroprotective recovery methods as previously described (*11*, *87*). Briefly, mice were anesthetized with 4% isoflurane and transcardially perfused with an ice-cold sucrose cutting solution containing: 234 mM sucrose, 2.5 mM KCl, 1.25 mM NaH_2_PO_4_, 10 mM MgSO_4_, 0.5 mM CaCl_2_, 26 mM NaHCO_3_, and 11 mM glucose (equilibrated with 95% O_2_ and 5% CO_2_, pH 7.4). Following perfusion, mice were decapitated, and brains were immediately immersed in the ice-cold cutting solution. Brains were sliced into 400 μm slices using a Leica VT1200 microtome (Leica Microsystems): coronal slices containing peri-injured somatosensory cortex were prepared for cortical studies, while horizontal slices containing the somatosensory ventrobasal thalamus (comprised of the VPL and VPM nuclei) and the nucleus reticularis thalami (nRT) were prepared for thalamic studies. 2-4 slices per animal were incubated at 32°C for 30 minutes, followed by 1 hour at 24–26°C, in artificial cerebrospinal fluid (aCSF) containing: 126 mM NaCl, 2.5 mM KCl, 1.25 mM NaH_2_PO_4_, 1 mM MgCl_2_, 2 mM CaCl_2_, 26 mM NaHCO_3_, and 10 mM glucose (300–310 mOsm, equilibrated with 95% O_2_ and 5% CO_2_, pH 7.4). For samples that were treated *in vivo* with IFNγ, IFNγ was added to the aCSF at 100 ng/mL (a concentration previously used for slice electrophysiology) (*62*, *64*) to maintain active IFNγ signaling *ex vivo* during recordings.

***Thalamic ex vivo electrophysiology***

Horizontal slices preserving intra-thalamic connectivity (described above) were transferred to an interface recording chamber maintained at 34°C. Slices were superfused at 2 mL/min with oxygenated aCSF supplemented with 0.3 mM glutamine (for metabolic support) (*11*, *87*).

*Recording Parameters:* Extracellular multi-unit activity was acquired using a linear 16-channel multi-electrode array (Neuronexus, Cat #A16x1-2mm-100-703) positioned across the nRT and VB regions (see Fig. 5F). Electrode placement was visually verified for each recording. Typically, electrodes #14-16 were positioned outside the VB area and were excluded from the analysis. Signals were amplified 10,000x and band-pass filtered between 100 Hz and 6 kHz using an RZ5 recording system (Tucker-Davis Technologies, SCR_006495).

*Stimulation Parameters:* Electrical stimuli to the internal capsule were delivered using bipolar tungsten microelectrodes (50–100 kΩ, FHC). Stimulation pulses were 100 μs in duration and 50 V in amplitude, delivered at 0.033 Hz (every 30 seconds) for a total of 10 trials per slice.(*11*, *87*)

*Quantification*: Extracellular spikes from multi-unit activity were detected with Burst script Spike2 (Version 8.21, Cambridge Electronic Design (CED), Cambridge, UK)). A “burst” of thalamic activity was defined as a cluster of ≥ 3 spikes occurring within 8ms intervals (with the interval between the first two spikes being ≤ 9 ms to officially signal a burst onset) (*88*–*91*). Evoked somatosensory thalamus bursts activity were analyzed by manually quantifying the number of evoked bursts in the first 3 seconds post stimulation (excluding the first 100 ms, where the direct response occurred) from the three the most responsive recording channels of each slice and averaged across 7 trials for each slice.

***Cortical local field potential recordings and laminar mapping***

Extracellular local field potentials (LFPs) were recorded from the perilesional cortex of TBI mice (or the corresponding somatosensory region in sham controls) in coronal brain slices as previously described (*83*).

*Electrode Configuration:* A linear 16-channel multi-electrode array with 100 μm inter-electrode spacing (Neuronexus, Cat #A16x1-2mm-100-703) was aligned normal to the cortical surface. The orientation of the probe was such that electrode #1 was always placed in layer 2/3 of the cortex and electrode #13 placed at the border of layer 6 and white matter. Electrode anatomical locations were as follows: #1–3 in layer 2/3; #4–5 in layer 4; and #6–12 in layers 5–6. Electrodes #13–16 were typically localized outside of the cortical area and were excluded from the analysis (see fig. S5F and G).

*Stimulation protocol and pharmacological dissection of evoked responses:* To evoke laminar synaptic responses, electrical stimulation was delivered in the underlying white matter (corpus callosum) with a concentric bipolar tungsten electrode (50–100 kΩ, FHC Inc.) as previously described (*83*). Electrical stimuli (100 μs pulse duration) were delivered every 8 seconds, sweeping through 10 escalating current intensities ranging from 10 μA to 600 μA. Each intensity was repeated 10 times per recording session.

*Data Acquisition:* Extracellular signals across all 16 channels were amplified 10,000x, band-pass filtered between 100 Hz and 6 kHz, digitized at 24.414 kHz, and recorded continuously using an RZ5 processor workstation (Tucker-Davis Technologies) (*83*).

*Pharmacological Isolation:* To biophysically isolate presynaptic and postsynaptic components, baseline aCSF recordings were followed by bath perfusion of glutamatergic receptor antagonists: the AMPA receptor antagonist 6,7-dinitroquinoxaline-2,3-dione (DNQX, 10 μM; Sigma-Aldrich) and the NMDA receptor antagonist 2-amino-5-phosphonopentanoic acid (APV, 125 μM; Sigma-Aldrich). Recordings were continued for 10 minutes while stimulus intensities were stepped from 50 μA to 500 μA. Slices were then washed in standard aCSF for 30 minutes. Finally, the voltage-gated sodium channel blocker tetrodotoxin (TTX, 1 μM, Sigma-Aldrich) was perfused for 10 minutes to eliminate all action potential propagation, and recordings were continued for 10 minutes with increasing stimuli up to 500 μA. The APV/DNQX-sensitive component was computationally designated as the post-synaptic response, and the residual TTX-sensitive component was defined as the pre-synaptic volley (*83*, *84*).

***Current source density computation***

To determine the precise laminar location, directional flow, and magnitude of evoked transmembrane currents, a current source density (CSD) analysis was performed (*83*, *84*). The CSD profile was computed by calculating the second spatial derivative of the recorded extracellular LFP voltage profiles along the cortical depth axis (z), using the following formula:

$$CSD(z) \approx-\sigma\frac{\phi(z+h) - 2\phi(z) + \phi(z-h)}{h^{2}}$$

where ϕ is the extracellular potential, h is the inter-electrode spacing (100 μm), and σ represents the macroscopic electrical conductivity of the tissue.

Inward net positive transmembrane currents (inward synaptic currents or active depolarization) were defined as current sinks (negative CSD deflections), while outward net currents were defined as current sources (positive CSD deflections). To visualize these spatiotemporal dynamics, color-mapped image plots were generated using custom MATLAB scripts (kindly provided by the Huguenard Lab) (*84*), where current sinks are represented in blue and current sources are represented in red. For more theory see Hodgkin and Huxley (*92*) and Mitzdorf (*93*).

Analysis of CSD data was performed as previously described (*83*). At each stimulus intensity, 10 recorded sweeps were averaged from each channel and each slice after excluding sweeps with occasional noise and small signal-to-noise ratio. In cases where LFPs had to be excluded, an assumed LFP was recovered by linearly interpolating between the LFPs from the two adjacent electrodes. The presynaptic response was measured between 0.3 and 6 ms after stimulation, and the postsynaptic response from 6 ms up to 200 ms (fig. S5J). We analyzed the amplitude of the negative and positive components of presynaptic and postsynaptic responses by measuring the baseline-to-peak value. Presynaptic and postsynaptic responses obtained from electrodes 1–3 were averaged and expressed as CSD of layers 2/3; presynaptic and postsynaptic responses obtained from electrodes 4, 5 were averaged and expressed as CSD of layer 4; and presynaptic and postsynaptic responses obtained from electrodes 6–12 were averaged and expressed as CSD of layers 5/6. Channels with noise were excluded from the analysis. For statistical analysis, n = 7–12 animals per group were included (1–2 slices per animal).

**Blood collection for flow cytometry**

Mice were deeply anesthetized with isoflurane. Blood was collected via cardiac puncture with an insulin syringe and thoroughly mixed with 0.3 mL of a 5 U/mL heparin sodium salt solution (Sigma) in 1X DPBS to prevent coagulation. Blood samples were then pelleted (500 x g, 5 min) and subjected to two rounds of red blood cell lysis (1X Pharm-Lyse solution; BD Biosciences) before antibody staining for flow cytometry. Final cell counts were normalized to the volume of blood collected from each mouse.

**Brain dissociation for flow cytometry**

Mice were deeply anesthetized with isoflurane and perfused transcardially with 10–15 mL of ice-cold 1X DPBS via the left ventricle (after blood collection, see above). The brain was dissected from the skull and three 1-mm-thick coronal sections encompassing the injury site were cut using a brain matrix. Perilesional cortex, ipsilateral hippocampus, and ipsilateral thalamus were microdissected under a dissection microscope and collected separately in 2 mL of iMED (isolation medium: 1X HBSS lacking calcium or magnesium, supplemented with 15 mM HEPES and 0.6% glucose). Brain samples were processed to isolate microglia and other immune cells as previously described (*82*). Briefly, tissues were mechanically dissociated in a 2 cm^3^ glass Dounce homogenizer in iMED on ice, filtered through a 70 μm cell strainer, and washed twice with 3 mL iMED. Samples were centrifuged at 220 x g for 10 min at 4°C and then resuspended in 22% Percoll solution (GE Healthcare) and overlaid with 1X DPBS. Samples were centrifuged at 950 x g for 20 min (acceleration 4, brake 0) and the supernatant layers were discarded to remove myelin and debris. Cell pellets were resuspended prior to antibody staining for flow cytometry.

**Staining for flow cytometry**

For myeloid and microglial panels, single cell suspensions were first incubated in FACS wash buffer (FWB: 1X DPBS (pH 7.4) with 3% (v/v) heat-inactivated FBS and 0.05% NaN_3_) containing 5% (v/v) normal rat serum and Fc Block (2.4G2, 1:250, BD Biosciences) for 20 min at 4°C. Cells were pelleted at 1200 rpm for 2 min and resuspended in FWB containing surface antibodies for 45 min at 4°C. Cells were washed and resuspended in FWB prior to analysis.

For lymphocyte panels involving transcription factor staining, single cell suspensions were first incubated with a viability dye (Fixable Viability Dye eFluor780, eBioscience), Fc Block (2.4G2, 1:100, BD Biosciences), and surface antibodies diluted in 1X DPBS for 45–60 min at 4°C. Cells were washed with FWB and then fixed and permeabilized overnight at 4°C using the FoxP3/transcription factor Staining Buffer Set (eBioscience). Cells were washed with 1X PermBuffer (eBioscience) and stained with intracellular antibodies for 60 min at 4°C. Cells were washed again with 1X PermBuffer and then resuspended in FWB prior to analysis. Any experiments that included anti-CD4 treated mice were stained with an anti-CD4 clone (RM4-5) different from the clone used for *in vivo* CD4 depletion (GK1.5) to avoid epitope masking.

For cytokine staining, cells were first stimulated *ex vivo* with PMA (phorbol 12-myristate 13-acetate) and ionomycin. Single cell suspensions were resuspended in complete RPMI (1X RPMI 1640 supplemented with 10% (v/v) heat-inactivated FBS, 1% (v/v) penicillin/streptomycin, 1% (v/v) Glutamax, 1 mM sodium pyruvate, 10 mM HEPES, 10 mM non-essential amino acids, and 55 μM β-mercaptoethanol) containing 1X Tonbo Cell Stimulation Cocktail (Cytek) and 1X Brefeldin A (BFA, eBioscience) for 3 hours in a 37°C / 5% CO_2_ incubator. Cells were washed with FWB and stained with a viability dye (Fixable Viability Dye eFluor780, eBioscience), Fc Block (2.4G2, 1:100, BD Biosciences), and surface antibodies diluted in 1X DPBS for 45–60 min at 4°C. Cells were then washed and fixed/permeabilized for 20 min at 4°C using the Cytofix/Cytoperm Fixation/Permeabilization Kit (BD Biosciences). Cells were washed with 1X PermBuffer (BD Biosciences) and stained with intracellular antibodies overnight at 4°C. Cells were washed again with 1X PermBuffer (BD Biosciences) and then resuspended in FWB prior to analysis.

All flow samples were stained in 50 μL in a 96-well V-bottom plate. See Table S2 for a list of antibodies used for flow cytometric staining. Prior to data collection, counting beads (CountBright Absolute Counting Beads, Invitrogen) were added to each sample and samples were filtered through a 40 μm cell strainer. Flow cytometric data was collected on a BD Fortessa (equipped with UV laser) and data analysis was performed using FlowJo software (BD).

**Immunofluorescence**

Mice were deeply anesthetized with isoflurane and perfused with 10–15 mL of ice-cold 1X PBS via the left ventricle, followed by 10 mL of 4% (w/v) paraformaldehyde (PFA) diluted in 1X PBS. Brains were dissected out and fixed overnight at 4°C in 4% PFA/PBS. Brains were then washed in 1X PBS and placed in 30% (w/v) sucrose for cryopreservation for a minimum of 2 days. Brains were either frozen in O.C.T. (Fisher) and stored at -80°C until sectioning on a cryostat (Leica) or flash-frozen on a freezing microtome (Microm). Brains were sectioned into 40 μm-thick sections, which were then stored in 1X DPBS containing 0.05% NaN_3_ at 4°C.

Brain sections were stained in 24 well plates and first blocked in 0.5 mL/well Block/Stain Solution (1X DPBS containing 5% (v/v) normal goat or horse serum and 0.4% (v/v) Triton X-100) for 1 hour at room temperature (RT). The normal serum used in the Block/Stain Solution corresponded to the species source of the secondary antibodies used (i.e., goat serum for goat-derived secondary antibodies or horse serum for donkey-derived secondary antibodies). Sections were then stained overnight at 4°C with primary antibodies diluted in 0.25 mL/well Block/Stain Solution on a plate shaker (~90rpm). Sections were washed four times with PBS-T (0.05% (v/v) Triton X-100 in 1X DPBS) for 5 min each at RT (~90 rpm) and then stained with secondary antibodies in 0.25 mL/well Block/Stain Solution for 1.5–2hrs at RT (~90 rpm). Sections were washed three times with PBS-T (5 min each at RT, ~90 rpm) before mounting in Fluoromount G with or without DAPI (Southern Biotech). Any antibodies directly conjugated to fluorophores were incubated with primary antibodies overnight and ensured to be compatible with subsequent secondary antibodies, if used. See Table S2 for a list of antibodies used for immunofluorescent staining.

**Confocal Imaging and Analysis**

Large tiled Z-stack images of stained sections were taken either on a Nikon A1R laser scanning confocal microscope (with 405, 488, 561, and 650 nm laser lines) or a Leica Stellaris 8 laser scanning confocal microscope (with a Leica LED3 390–690 nm white light laser). On the Nikon A1R, large tiled images of hemi-brains were collected using a 16X/0.8 NA Plan Apo water-immersion objective with 512x512 resolution, 1 frame/s scanning speed, 2x line averaging, and 4μm Z-steps. Higher magnification tiled Z-stack images of thalami were collected using a 25X/1.1 NA Plan Apo water-immersion objective with 1024x1024 resolution, 0.5 frame/s scanning speed, 2x line averaging, and 2 μm Z-steps. On the Leica Stellaris, all images were collected using a 20X/0.75 NA CS2 air objective, 512x512 resolution, bidirectional scanning, 400 scan speed, 2x line averaging, and 3μm Z-steps.

High magnification images were taken on a Zeiss LSM800 laser scanning confocal microscope using a 40X/1.4 DIC oil-immersion objective, 1024x1024 resolution, bidirectional scanning, 4 scan speed, and 4x line averaging. Each image taken was a single plane field of view.

Images were rendered into three dimensions and quantitatively analyzed using Bitplane Imaris v9.5.1 software (Andor Technology PLC). 3D reconstructions of each channel were generated and quantified using the Imaris surface function, thresholded on signal intensity, volume, and/or DAPI-intensity, as previously described (*28*, *81*, *94*). Thalamic subregions were manually traced in Imaris based on NeuN staining while referencing the adult coronal Allen Brain Atlas (*95*), and cell counts per region were normalized to the cross-sectional area of each subregion. Large tiled Z-stack images are all displayed as maximum intensity projections. Two sections per mouse were imaged and averaged for analysis.

**Brain tissue homogenization and cytokine measurement**

Mice were deeply anesthetized with isoflurane and perfused with 10-15 mL of ice-cold 1X DPBS via the left ventricle. The brain was dissected from the skull and three 1mm coronal sections were cut using a brain matrix. Perilesional cortex, ipsilateral hippocampus, and ipsilateral thalamus were micro-dissected under a dissection microscope from these sections, snap frozen on dry ice, and stored at -80°C. Tissue samples were thawed on ice, weighed, and transferred to a 2 mL tube pre-filled with 3.0 mm high-impact zirconium beads (Benchmark Scientific). 1X RIPA Buffer (Cell Signaling Technology) supplemented with 1 mM PMSF (Cell Signaling Technology) was added to each tube at a ratio of 1 mL per 100 mg of tissue per manufacturer’s recommendations. Samples were lysed and homogenized on a Precellys Evolution Touch Homogenizer (Bertin Technologies) on the standard mouse setting (2x30s at 6500 rpm) at 4°C. Crude lysates were centrifuged at 10,000 x g for 10min at 4°C and the clarified supernatant was collected. Total protein content of each homogenate was quantified using the Pierce BCA Protein Assay Kit (Thermo Scientific) and homogenates were stored at -80°C. Homogenates were then normalized to 2 mg/mL total protein and sent to Eve Technologies (Calgary, Canada) on dry ice. Eve Technologies performed Luminex multiplex cytokine measurement to quantify cytokine concentrations in the homogenates using the Mouse High-Sensitivity T Cell 18-Plex Discovery Assay (MDHSTC18) Array. Total cytokine quantities were calculated from measured cytokine concentrations and normalized to the weight of the original tissue sample.

**CD11b^+^ cell isolation and single cell RNA sequencing (scRNAseq)**

For microglial isolation for downstream RNA-sequencing, cells were isolated as described previously (*82*). Briefly, the right (ipsilateral) thalami from four mice per condition (two male and two female) were dissected on ice, pooled, and mechanically dissociated using a glass tissue homogenizer in isolation medium (1X HBSS, 15 mM HEPES, 0.6% glucose, 1 mM EDTA). Cells were filtered through 70 µm cell strainers and then centrifuged at 300 x g for 10 minutes at 4°C before being resuspended in 22% Percoll (GE Healthcare) and centrifuged at 950 x g for 20 minutes (acceleration 4, brake 0) in order to remove myelin and cellular debris. Pelleted cells were then resuspended in staining buffer (1X PBS, 0.5% BSA, 2 mM EDTA) and incubated with CD11b MACS beads (Miltenyi Biotech, 1:50) for 15 minutes at 4°C. Cells were washed with staining buffer, pelleted at 300 x g for 5 minutes at 4°C, and reconstituted in 500 µL staining buffer. CD11b^+^ myeloid cells were isolated as described in the manual for MACS LS columns (Miiltenyi Biotech) and collected in a staining buffer without EDTA, pelleted at 300 x g for 5 minutes at 4°C, and counted on a hemocytometer. 15,000–20,000 cells were diluted in 30 µL in a BSA-coated plate for 10x Genomics sequencing.

Approximately 40,000 cells were loaded into each well of Chromium Chip G (v3.1) on the Chromium X, libraries were prepared in-house as described in the 10x Manual, and libraries were sequenced on three lanes of the NovaSeq SP100 at the UCSF Center for Advanced Technologies (CAT) core.

**scRNAseq data analysis**

Sequenced samples were processed using the Cell Ranger 2.1 pipeline (built on the STAR aligner) (*96*) and aligned to the GRCm38 (mm10) mouse reference genome. Clustering and differential expression analysis were conducted using Seurat version 3.1.4 (Table S3). Sequencing scripts can be found at https://github.com/lcdorman/Paz_MG_TBI, and original data can be found on GEO (accession number to be provided upon acceptance).

Cells outside of the thresholds listed in Table S4 were excluded from downstream analysis. Cells were identified as “female” or “male” based on their expressions of the genes *Xist, Tsix, Ddx3y, and Eif2s3y*; any cells expressing at least one count of *Xist* or *Tsix* and no counts of *Ddx3y/Eif2s3y* were labeled female, while all others were labeled male. Counts were then normalized using sctransform, regressing out percent mitochondrial RNA and total genes per cell. The top 2000 most variable genes were used to calculate 50 principal components, and the top 30 PCs were used for nearest neighbor, UMAP, and cluster calculations (resolution 1). Contaminating cell types were identified through expression of *Slc1a3* and *Mbp*, and all cells with normalized expression of *Mbp* > 2 were removed. Microglia were then re-normalized and clustered as described above, with a resolution of 0.5. Two sets of most closely related homeostatic clusters (0/1/5/6/7 and 3/9) were combined due to <10 upregulated genes (log fold change > 0.2, adjusted p-value < 10^-8^).

Differential gene expression between clusters was calculated using the MAST test in Seurat. Volcano plots were generated using the EnhancedVolcano package in R, with gene labels chosen from the list of differentially expressed genes. Cutoffs were set at log2 fold change of 0.37 (30% increase) and adjusted p-value < 10^-25^.

Bar plots depicting relative cluster frequency were created using ggplot2 in R. Cell counts for each cluster were first normalized to the total number of cells recovered from the corresponding sample to account for differences in sample size. For each cluster, the normalized counts were then converted to percentages by dividing the normalized count from each sample by the sum of normalized counts across all samples within that cluster and multiplying by 100.

**Stereotactic i.c.v. injection, microglial isolation, and bulk RNA sequencing**

Brain injections were performed with a Kopf stereotaxic apparatus (David Kopf, Tujunga, CA) and a microdispensing pump (World Precision Instruments) holding a Hamilton Syringe (model 701 RN, 10 microliter) with a beveled glass needle (~50 micron outer diameter). For intraventricular (i.c.v.) injections into juveniles (P9), mice were anesthetized with 1.5% isoflurane at an oxygen flow rate of 1L/min, head-fixed with a stereotaxic frame (for juveniles, size P11), and treated with ophthalmic eye ointment. Fur was shaved and the incision site was sterilized with 70% ethanol and Betadine prior to surgical procedures. Subcutaneous 0.5% lidocaine was administered at the incision site and lack of reflex response was checked. Body temperature was maintained throughout surgery using a heating pad. After incision, a hole was drilled in the skull and 500 nL of recombinant mouse IFNγ (diluted to 0.2 mg/ml in PBS, Gibco) or PBS was injected (from lambda: 3 mm AP, 1.5 mm ML, -2 mm DV) at a rate of 250 nL/min. The needle was held in place for 5 minutes to allow diffusion and then slowly removed. The incision was closed and mice were allowed to fully recover with heat. Buprenorphine (Henry Schein Animal Health) was administered (0.1 mg/kg) according to approved protocols (briefly, mice were dosed prior to surgery, 4–8h later, and the next morning if needed by intraperitoneal injection).

Approximately 22h later, mice were euthanized and microglia were isolated from cortex, as described previously (*82*). 2 females and 1 male mice were used per condition, aged P10 at collection. Cells were stained with CD45–FITC, CD11b–PE, and Ly6C–APC (See Table S2 for details). CD45^lo^ CD11b^+^ Ly6C^-^ microglia were sorted on a BD Aria III sorter into RLT plus buffer (QIAGEN). RNA was isolated from 60,000–100,000 microglia per mouse with the RNeasy® Plus Micro kit (Qiagen). Quality and concentration were determined with the Agilent RNA 6000 Pico kit on a Bioanalyzer (Agilent). All samples had an RNA Integrity Number (RIN) >7. cDNA and libraries were made using the Lexogen QuantSeq 3’ mRNA-seq FWD library prep kit and quality was assessed by Agilent High Sensitivity DNA kit on a Bioanalyzer (Agilent). Pooled libraries were RNA sequenced on an Illumina HiSeq 4000 single-end for 65 cycles (SE65) yielding 50–70 million reads per sample. Quality of reads was evaluated using FastQC (http://www.bioinformatics.babraham.ac.uk/projects/fastqc), all samples passed quality control, and reads were aligned to mm10 (GRCm38; retrieved from Ensembl, version September 2017) using STAR (version 2.5.4b) (*96*) with ‘–outFilterMultimapNmax 1’ to only keep reads that map one time to the reference genome. Mapped reads were counted using HTSeq (version 0.9.0)(*97*) and DESeq2 package (version 1.24.0) (*98*) was used to normalize the raw counts and perform differential gene expression analysis.

**Supplementary Figures:**

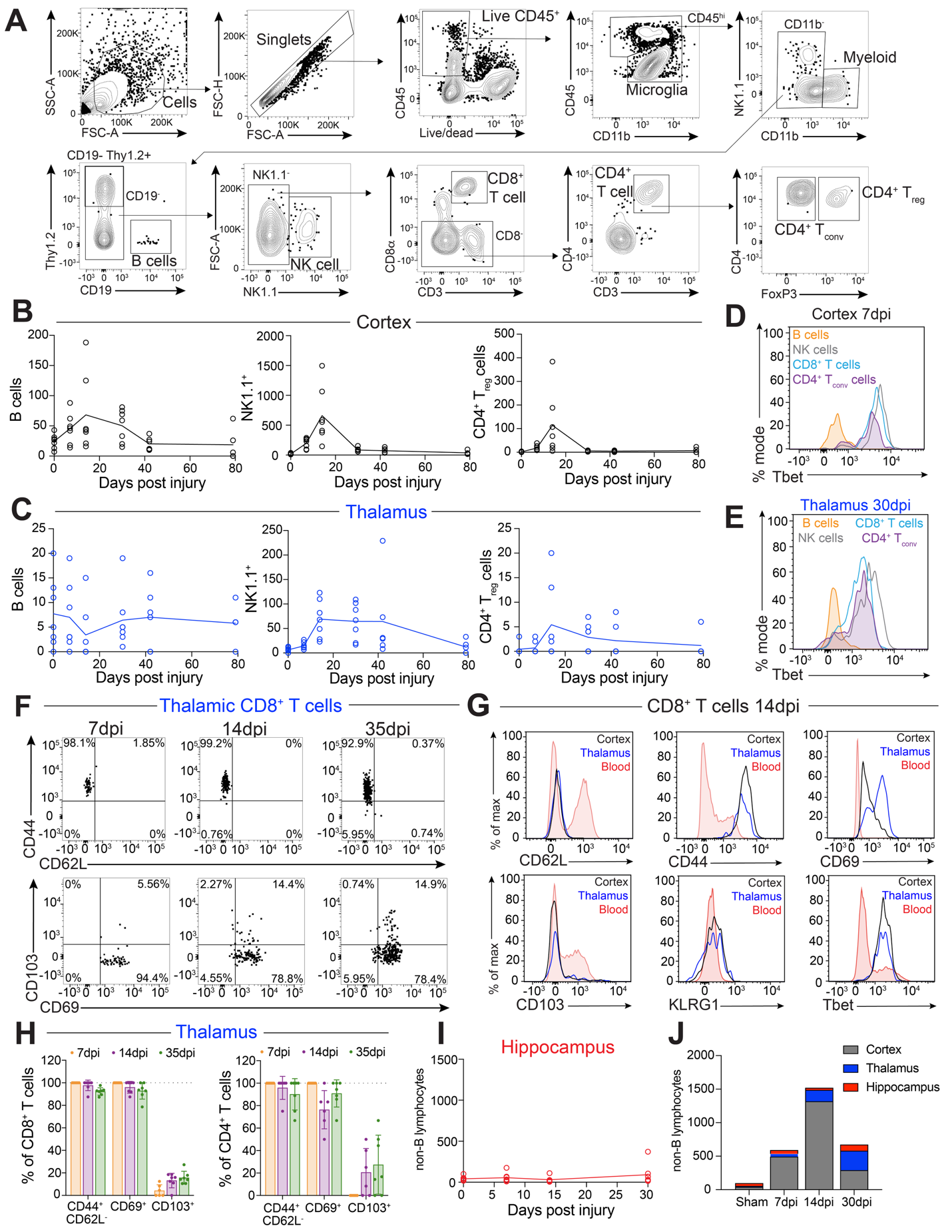

**Fig. S1. Characterization of immune infiltrates after TBI, kinetic profiles and surface phenotyping, related to Fig. 1.**

**(A)** Flow cytometric gating strategy used to identify brain-infiltrating lymphocyte populations, with representative plots derived from a perilesional cortical sample at 7 dpi. Related to **Fig. 1**. Representative flow plots are from a cortical sample at 7 dpi.

**(B and C)** Temporal dynamics showing absolute numbers of B cells (CD19^+^), NK/ILC1s (NK1.1^+^), and regulatory CD4^+^ FoxP3^+^ T cells (T_regs_) isolated from the perilesional cortex (B) and the ipsilateral thalamus (D).

**(D and E)** Representative flow cytometric histograms showing intracellular Tbet expression across indicated lymphoid subsets within the cortex at 7 dpi (C) and the thalamus at 30dpi (E) validating cell type-specific staining.

**(F)** Concatenated flow cytometric plots of thalamic CD8^+^ T cells across the post-injury timecourse, tracking surface expression profiles of activation and tissue-residency markers (CD44, CD62L, CD69, and CD103).

**(G)** Representative flow cytometric histograms comparing expression of CD62L, CD44, CD69, CD103, KLRG1, and Tbet among CD8^+^ T cells isolated from the cortex, thalamus, and peripheral blood at 14 dpi.

**(H)** Relative proportions of phenotypic marker-positive (CD44^+^ CD62^+^, CD69^+^, and CD103^+^) subsets within the thalamic CD8^+^ T cell pool (left) and CD4^+^ T cell pool (right) across the longitudinal timecourse.

**(I)** Absolute quantification of infiltrating non-B lymphocytes (CD45^hi^ CD11b^−^ CD19^−^ Thy1.2^+^) isolated from the ipsilateral hippocampus over time.

**(J)** Spatiotemporal comparison of total infiltrating non-B lymphocyte kinetics mapped across the perilesional cortex, ipsilateral hippocampus, and ipsilateral thalamus after TBI. Data represents means.

**Data and Sample Sizes:** Main panels represent mean values only for (B, C, and J) to emphasize kinetic trends, and mean ± SD for (H). Individual data points represent unique mice (biological replicates).

- **Sample Sizes:** For cortical, thalamic, and hippocampal longitudinal mapping (B, C, I, and J), sample sizes per post-injury timepoint are: 7 dpi n = 6; 14 dpi n = 7; 35 dpi n = 7. For thalamic phenotyping and kinetics (F and H), n = 7 mice per individual timepoint.
- **Statistics:** Evaluated via mixed-effects model (REML) for (B and D): For (B), p = 0.0360 (B cells), p < 0.0001 (NK1.1^+^), p = 0.0061 (CD4^+^ T_regs_). (D), p = 0.7766 (B cells), p = 0.0093 (NK1.1^+^), and p = 0.2259 (CD4^+^ T_regs_).

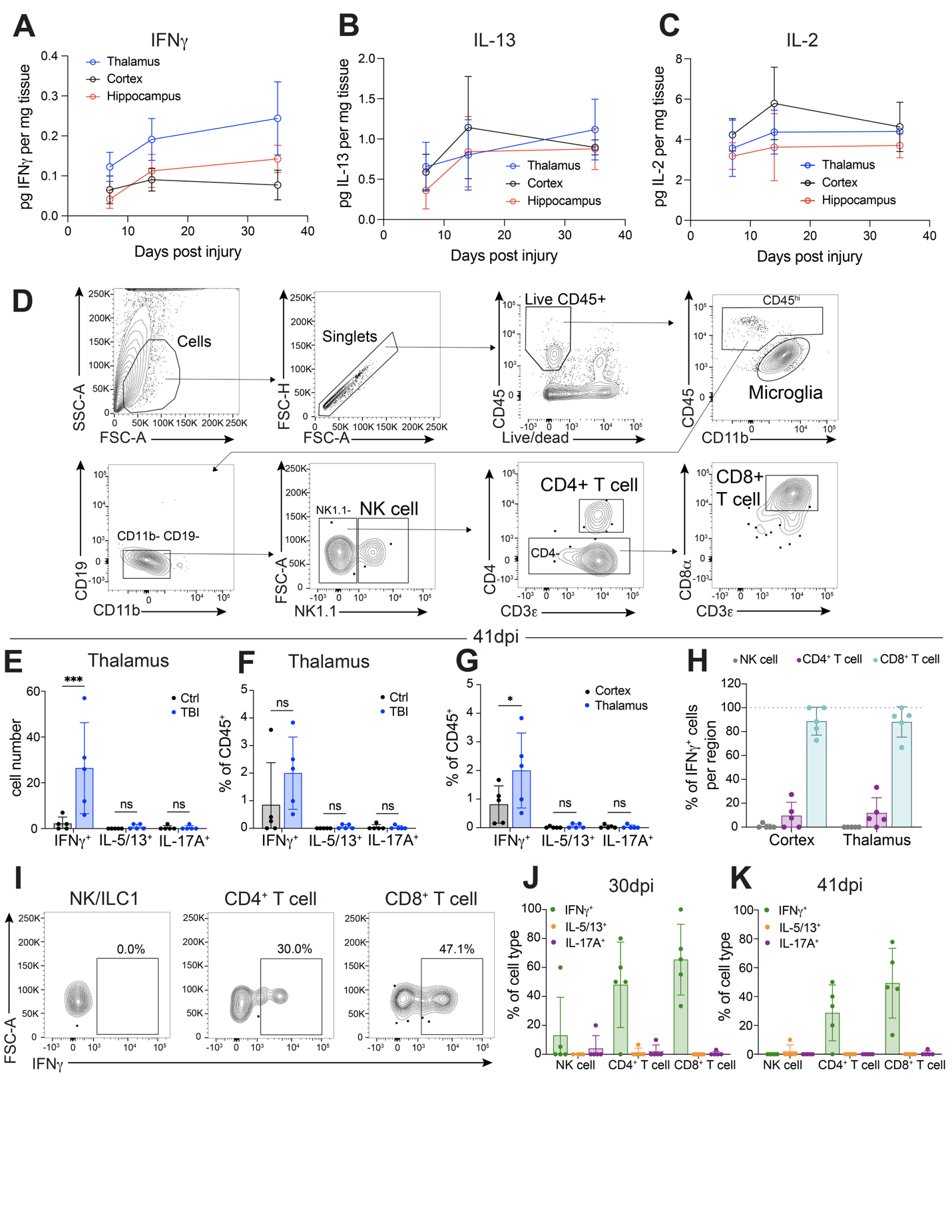

**Fig. S2. Spatial cytokine profiling and immunophenotypic characterization of cytokine-producing lymphoid subsets, related to Fig. 2.**

**(A to C)** Spatial tracking of IFNγ (A), IL-13 (B), and IL-2 (C) concentrations compared across the perilesional cortex, ipsilateral hippocampus, and ipsilateral thalamus over a longitudinal post-injury timecourse (7, 14, and 35 dpi) measured via multiplex bead immunoassay on micro-dissected tissue homogenates. Cytokine values are normalized to total tissue weight (mg).

**(D)** Flow cytometric gating strategy used to identify brain-infiltrating lymphocytes for intracellular cytokine quantification following *ex vivo* stimulation, with representative plots derived from cortical samples at 41 dpi. Related to Fig. 2).

**(E and F)** Total cell numbers (E) and relative frequencies within total CD45^+^ brain-infiltrating leukocytes (F) of IFNγ^+^, IL-5/13^+^, and IL-17A^+^ cells in the thalamus of sham versus TBI mice at 41 dpi.

**(G)** Relative proportion within total CD45^+^ infiltrating leukocytes of IFNγ^+^, IL-5/13^+^, and IL-17A^+^ cells in the perilesional cortex versus ipsilateral thalamus at 41 dpi.

**(H)** Immunophenotypic composition of the total reactive IFNγ^+^ pool at 41 dpi, comparing the relative frequency of constituent lymphoid subsets between the perilesional cortex and ipsilateral thalamus.

**(I)** Representative flow cytometric plots displaying IFNγ expression within thalamic NK/ILC1s, CD4^+^ T cells, and CD8^+^ T cells at 41 dpi.

**(J and K)** Proportional frequencies of cytokine-positive (IFNγ^+^, IL-5/13^+^, and IL-17A^+^) cells within lymphoid lineages evaluated at 30 dpi (F) and 41 dpi (G).

**Data and statistics:** Bar graphs and data points represent mean ± SD from individual mice (biological replicates).

- **Sample sizes:** For regional cytokine mapping (A to C), 7 dpi (cortex n = 6, hippocampus n = 3, thalamus n = 5); 14 dpi (cortex n = 6, hippocampus n = 4, thalamus n = 5); 35 dpi (cortex n = 7, hippocampus n = 5, thalamus n = 6). For (F, G, J, and K), TBI n = 5 mice. For (H and I), sham n = 4; TBI n = 5.
- **Statistical analysis:** Evaluated via two-way ANOVAs with Šidák’s multiple comparisons test for (H, I, and J). Statistics: ns = not significant, *p < 0.05, ***p <0.001.

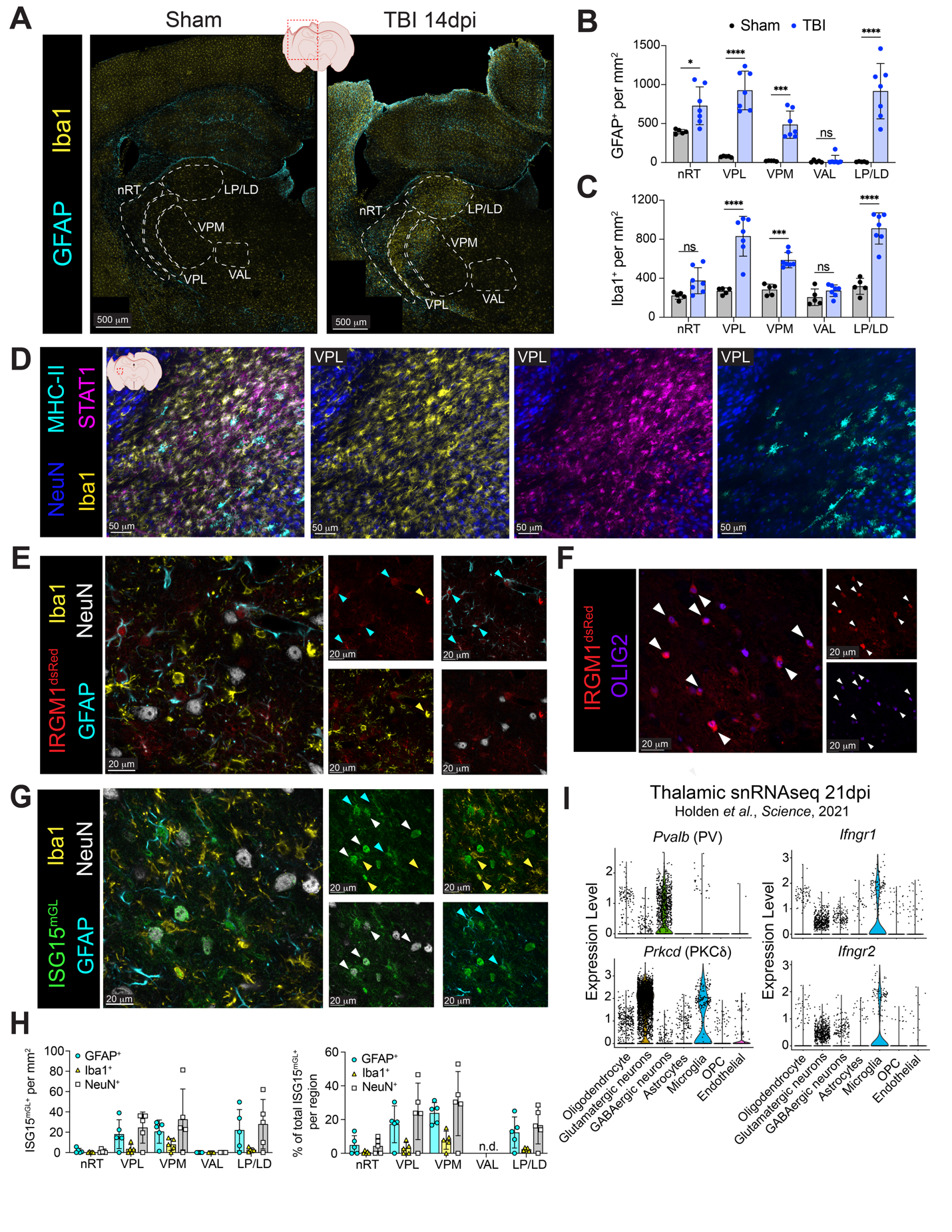

**Fig. S3. Glial reactivity profiles and cellular distribution of interferon-responsive reporter expression within the thalamus, related to Fig. 3.**

**(A)** Representative confocal images of the ipsilateral brain hemisphere from sham and TBI mice at 14 dpi, displaying immunofluorescence for GFAP (astrocytes) and Iba1 (microglia/ macrophages). Thalamic subregions are delineated (dashed white lines) based on NeuN counterstaining (not shown).

**(B and C)** Spatial quantification of GFAP^+^ **(B)** and Iba1^+^ **(C)** cell density across designated thalamic subregions.

**(D)** Representative confocal image of the ipsilateral ventroposterolateral (VPL) thalamus after TBI (14 dpi) depicting immunofluorescence for NeuN, Iba1, MHC-II, and STAT1 immunofluorescence.

**(E)** Representative high-magnification confocal images of the ipsilateral thalamus from *Irgm1^dsRed+^* reporter mice at 14 dpi depicting IRGM1^dsRed^, Iba1, GFAP, and NeuN immunofluorescence. Cyan arrowheads indicate IRGM1^dsRed+^ GFAP^+^ astrocytes, yellow arrowhead indicates IRGM1^dsRed+^ Iba1^+^ microglia/macrophages.

**(F)** Representative high-magnification confocal image in the ipsilateral thalamus of a *Irgm1^dsRed+^* reporter mouse at 14 dpi depicting IRGM1^dsRed^ and OLIG2 immunofluorescence. White arrowheads indicate IRGM1^dsRed+^ OLIG2^+^ oligodendrocyte-lineage cells.

**(G)** Representative high-magnification confocal image of the ipsilateral thalamus from *Isg15^mGL/+^* reporter mice at 14 dpi depicting ISG15^mGL^, GFAP, Iba1, and NeuN staining. Respective cellular sources of ISG15^mGL^ are highlighted by cyan (astrocytes), yellow (microglia/macrophages), and white (neurons) arrowheads.

**(H)** Absolute cell density (left) and relative lineage proportions (right) of ISG15^mGL+^-expressing astrocytes, microglia/macrophages, and neurons across distinct thalamic subregions.

**(I)** Single-nucleus RNA sequencing (snRNA-seq) expression profiles of *Pvalb* (PV), *Prkcd* (PKCδ), *Ifngr1*, and *Ifngr2* across established thalamic cell clusters, reanalyzed from a public post-TBI dataset at 21 dpi (Holden *et al.*, *Science*, 2021). Cell cluster assignments are as depicted in the original publication (*10*).

**Data and statistics:** Bar graphs and data points represent mean ± SD from individual mice (biological replicates), except in (I) where data points are individual sequenced cells. Thalamic subregions: reticular nucleus (nRT), ventral posterolateral (VPL), ventral posteromedial (VPM), ventral anterolateral (VAL), and lateral posterior/lateral dorsal (LP/LD). n.d. = none detected.

- **Sample size:** For (B and C), sham n = 5; TBI n = 7 mice. For (H), TBI n = 5 mice.
- **Statistics:** For (B and C), two-way ANOVA with Šidák’s multiple comparisons test. ns = non-significant, *p < 0.05, ***p < 0.001, ****p < 0.0001.

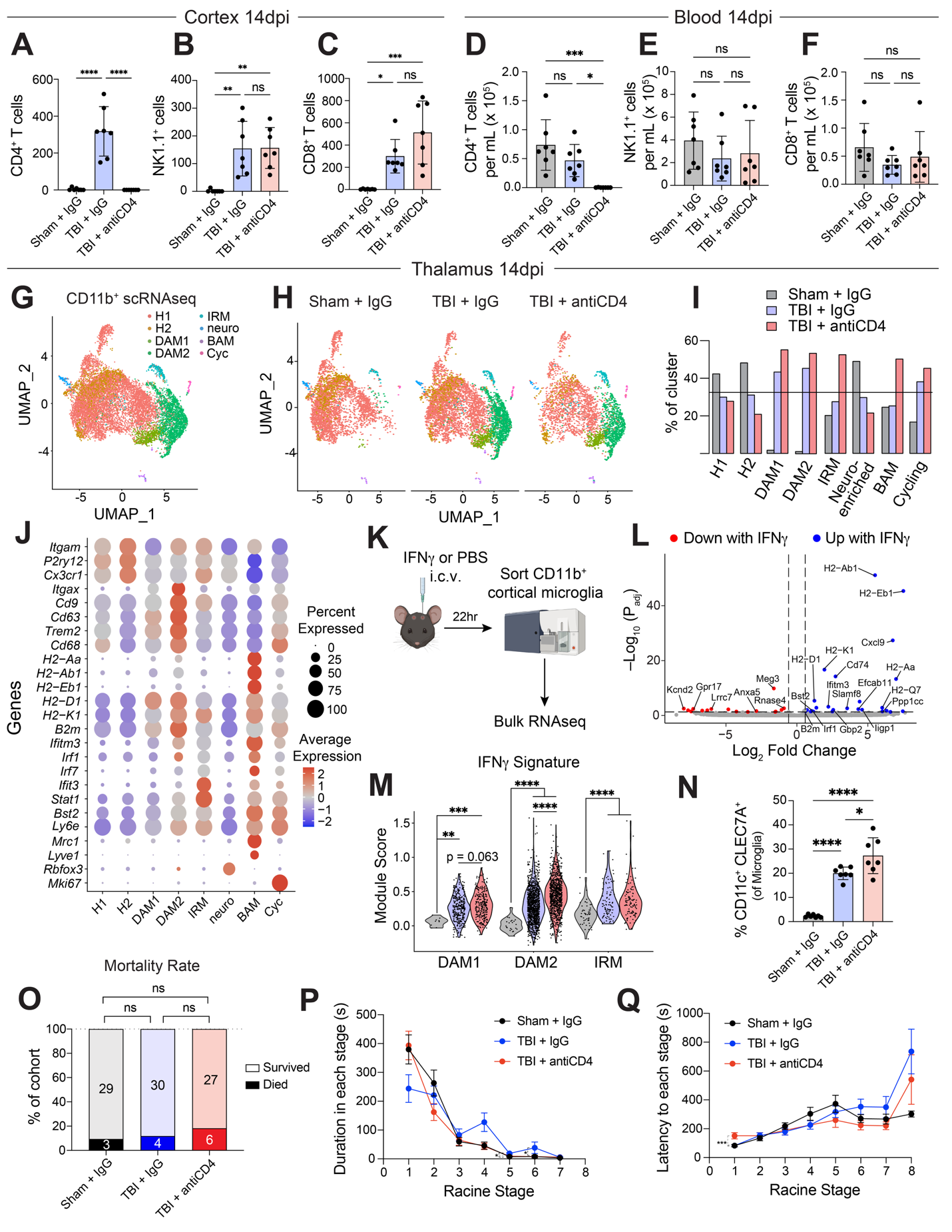

**Fig. S4. Extended phenotypic, transcriptomic, and flow cytometric characterization of CD4^+^ T cell depletion cohorts, related to Fig. 4.**

**(A to C)** Absolute number of infiltrating perilesional cortical CD4^+^ T cells (A), NK cells (NK1.1^+^) (B), and CD8^+^ T cells (C) at 14 dpi.

**(D to F)** Concentration of circulating CD4^+^ T cells (D), NK cells (NK1.1^+^) (E), and CD8^+^ T cells (F) in the blood.

**(G)** Unsupervised clustering visualization of scRNA-seq profiles from CD11b^+^ microglia and myeloid cells pooled across groups.

**(H)** Unsupervised clustering visualization of scRNA-seq clusters stratified by experimental group, downsampled to display 2,500 cells per condition.

**(I)** Relative frequency of each cluster (out of total myeloid cells) across experimental conditions. The horizontal dashed line (33%) denotes the theoretical baseline for uniform sample distribution across the three pooled groups.

**(J)** Dot plot illustrating expression of selected marker transcripts across myeloid clusters (as in (G)); circle size denotes the percentage of cluster cells expressing the gene, and heatmap color intensity reflects relative average expression.

**(K)** Experimental paradigm illustrating intracerebroventricular (i.c.v.) administration of recombinant IFNγ (or PBS vehicle) followed by isolation and fluorescence-activated cell sorting (FACS) of cortical microglia (CD45^lo^ CD11b^+^ Ly6C^–^) 24 hours post-injection for bulk RNA- sequencing.

**(L)** Volcano plot depicting differentially expressed genes (DEGs) in cortical microglia 24 hours after *in vivo* IFNγ exposure, compared to vehicle controls (as shown in (K)). Highlighted data points meet significance thresholds of adjusted p < 0.05 and |fold-change| > 1.5.

**(M)** IFNγ transcriptional signature module scores calculated per individual cell across disease-associated microglia (DAM1, DAM2) and IFN-responsive (IRM) myeloid clusters, stratified by treatment group (see fig. S4 J, K, and N).

**(N)** Proportion of CD11c^+^ CLEC7A^+^ disease-associated microglia (DAMs) within the total thalamic microglia pool.

**(O to Q)** Behavioral seizure metrics following PTZ challenge, displaying the proportion of each cohort experiencing post-challenge mortality within 20 minutes (A), cumulative duration spent within individual Racine severity stages (B), and latency to the initial onset of each stage (C).

**Data and Statistics:** Graphs represent mean ± SEM for (B and C) and mean ± SD for (J to O). Individual data points represent unique biological replicates (independent mice), except in (F and H) where individual data points represent single sequenced cells.

- **Sample size:** For (A to F, and N), n = 7 per group. For (O to Q), sham+IgG n = 32, TBI+IgG n = 34, TBI+anti-CD4 n = 33.
- **Statistical Analysis:** Evaluated via one-way ANOVA with Tukey’s post-hoc test for (A to F, and N), Kruskall-Wallis test with Dunn’s multiple comparisons test for (M, P and Q), and Fisher’s exact test for (O). Statistics: ns = not significant, *p < 0.05, **p < 0.01, ***p < 0.001, ****p < 0.0001.

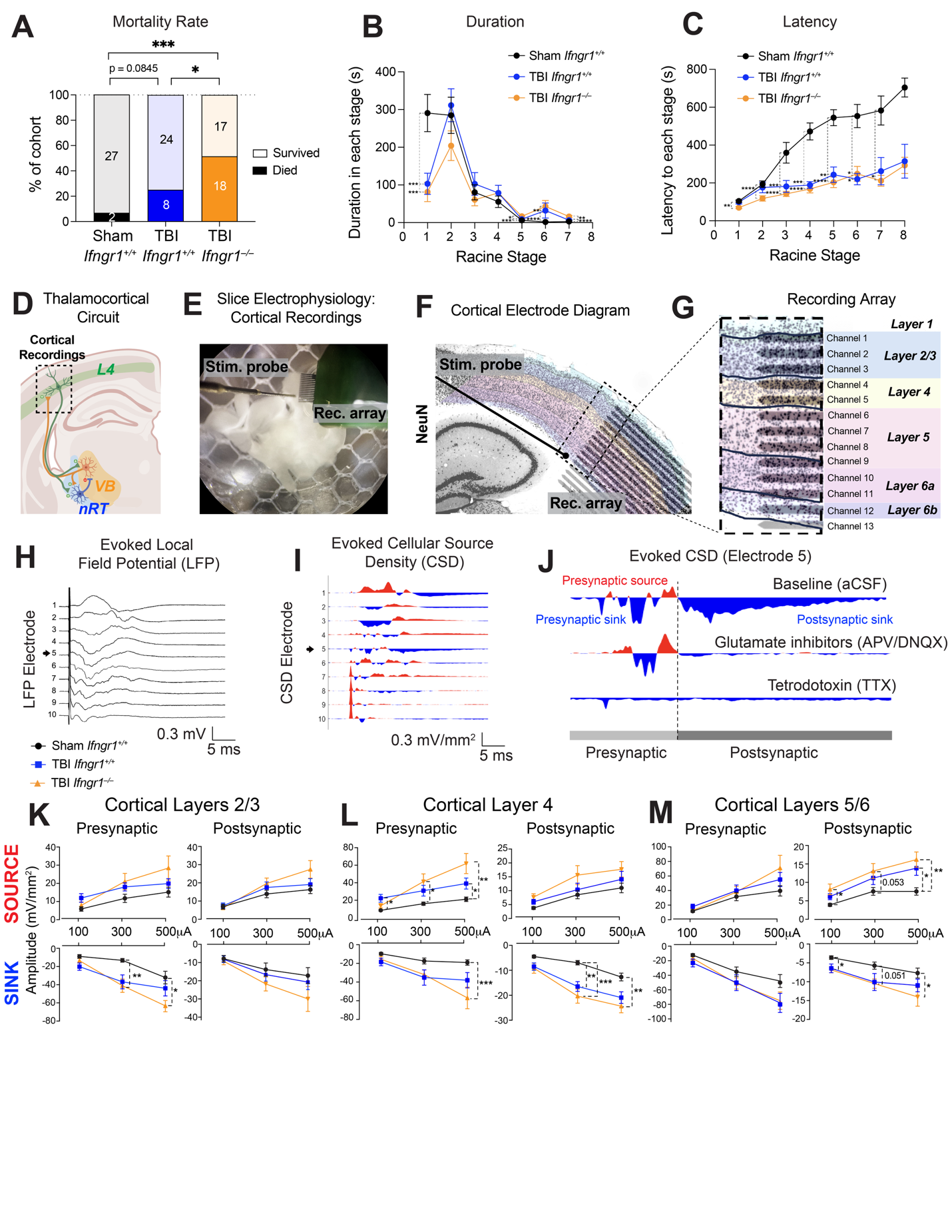

**Fig. S5. Behavioral seizure kinetics and laminar current source density analysis of cortical circuits in global Ifngr1 knockouts, related to Fig. 5.**

**(A to C)** Behavioral seizure metrics following PTZ challenge, displaying the proportion of each cohort experiencing post-challenge mortality within 20 minutes (A), cumulative duration spent within individual Racine severity stages (B), and latency to the initial onset of each stage (C).

**(D)** Diagram of thalamocortical circuit highlighting relevant circuitry for cortical recordings.

**(E and F)** Photo (E) and schematic (F) of cortical laminar electrophysiological array recordings and stimulation of the white matter (corpus callosum).

**(G)** Schematic mapping individual recording electrode channels to specific anatomical cortical layers.

**(H to J)** Representative evoked cortical electrophysiological recordings, displaying a characteristic local field potential (LFP) trace (H), corresponding current source density (CSD) profiles (I), and control validation CSD waveforms from a representative channel under baseline artificial cerebrospinal fluid (aCSF), glutamatergic receptor antagonism (APV/DNQX), or action potential-dependent neurotransmission blockade (tetrodotoxin; TTX) to isolate pre- versus post-synaptic components (J).

**(K to M)** Quantitative laminar CSD analysis tracking presynaptic and postsynaptic sink and source amplitudes across cortical layers 2/3 (J), layer 4 (K), and layers 5/6 (L) among the indicated experimental groups.

**Data and Statistics:** Graphs represent mean ± SEM. Individual data points represent individual brain slices in (K to M).

- **Sample size:** For (A to C), sham *Ifngr1^+/+^* n = 29 mice; TBI *Ifngr1^+/+^* n = 32 mice; TBI *Ifngr1^–/–^* n = 35 mice. For (K to M), sham *Ifngr1^+/+^* n = 16 slices from n = 12 mice ; TBI *Ifngr1^+/+^* n = 13 slices from n = 7 mice ; TBI *Ifngr1^–/–^* n = 11 slices from n = 6 mice.
- **Statistical Analysis:** Evaluated via Fisher’s exact test for (A) and Kruskal-Wallis test with Dunn’s multiple comparisons test for (B, C, and K to M). Statistics: ns = not significant, *p < 0.05, **p < 0.01, ***p < 0.001, ****p < 0.0001.

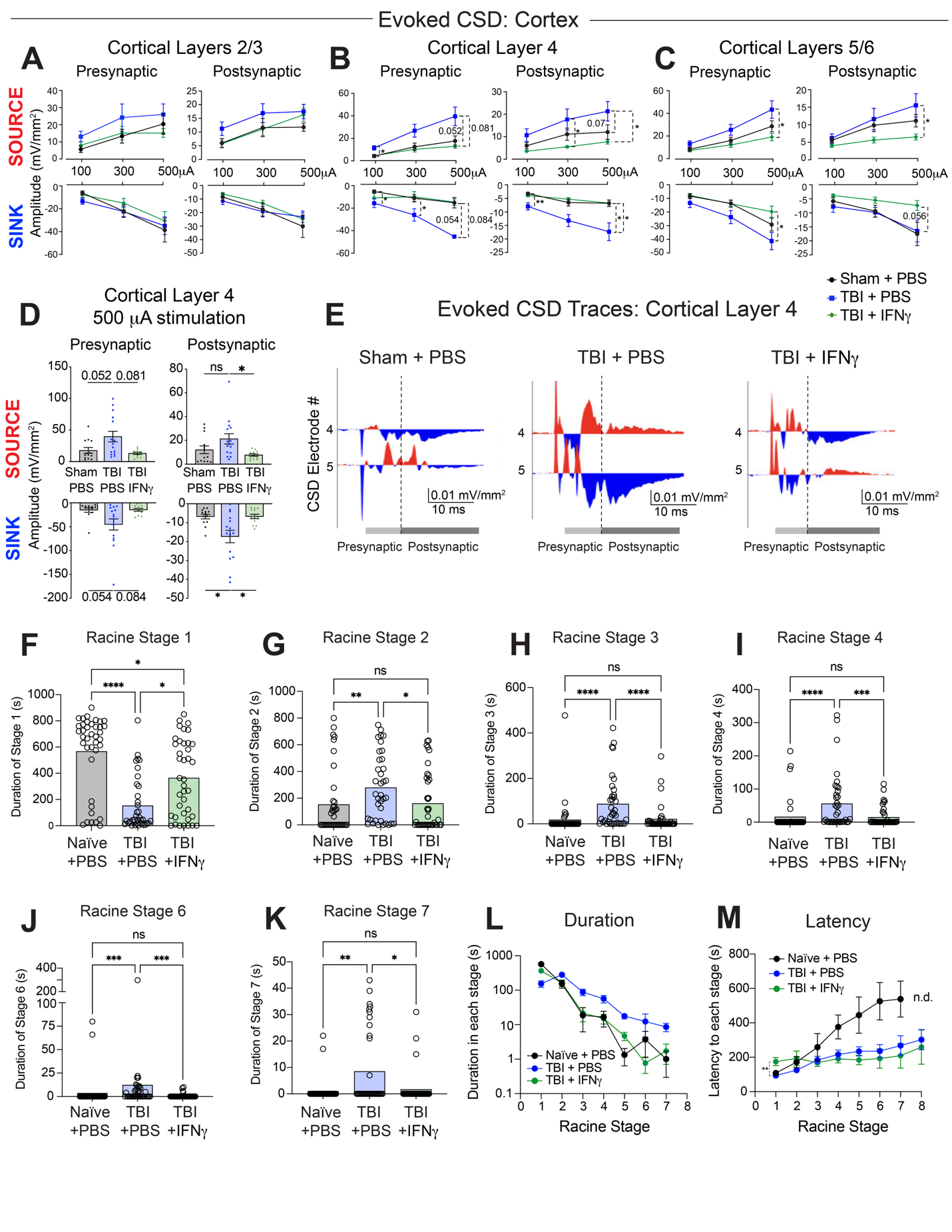

**Fig. S6. Cortical circuit current source density analysis and granular behavioral seizure profiles following therapeutic IFNγ administration, related to Fig. 6.**

**(A to C)** Laminar current source density (CSD) analysis tracking presynaptic and postsynaptic sink and source amplitudes across cortical layers 2/3 (A), layer 4 (B), and layers 5/6 (C) among the indicated treatment groups.

**(D and E)** Targeted CSD profiling within cortical layer 4 under a fixed 500 µA stimulation intensity, showing quantitative slice-level amplitudes (D) and corresponding representative CSD waveforms across recording channels (electrodes 4 and 5) (E).

**(F to K)** Classification of behavioral seizures following PTZ challenge at 5 weeks post-injury (wpi), detailing the individual duration spent within Racine Stage 1 (F), Stage 2 (G), Stage 3 (H), Stage 4 (I), Stage 6 (J), and Stage 7 (K).

**(L to M)** Summary metrics of behavioral seizure phenotypes, tracking the cumulative duration spent within (L) and latency to the initial onset of (M) each individual Racine stage.

**Data and Statistics:** Graphs represent mean ± SEM except for (F to K) which show means. Individual data points represent individual brain slices in (D) or unique biological replicates (independent mice) in (F to K).

- **Sample size:** For (A to D), naïve + PBS n = 12–14 slice from n = 7 mice ; TBI + PBS n = 12 slices from n = 11 mice ; TBI + IFNγ n = 12 slices from n = 7 mice. For (J to M), naïve + PBS n = 39 mice, TBI + PBS n = 37 mice, TBI+ IFNγ n = 39 mice.
- **Statistical Analysis:** Evaluated via Kruskal-Wallis test with Dunn’s multiple comparisons test within each stimulation current (A to D) or Racine stage (F to M). Statistics: ns = not significant, *p < 0.05, **p < 0.01, ***p < 0.001, ****p < 0.0001. n.d. = no data.

**Supplementary Tables:**

**Table S1. Modified Racine Scale used for classification and quantification of behavioral seizures.**

| **Racine Stage** | **Associated Behavioral Manifestations** |
| --- | --- |
| 0 | Baseline |
| 1 | Sudden behavioral arrest, immobility |
| 2 | Myoclonic jerks, mild |
| 3 | Myoclonic jerks, moderate to severe, with stiffened/extended tail |
| 4 | Partial body clonus, forelimb or hindlimb |
| 5 | Generalized tonic-clonic (GTC) seizure, with loss of posture/control |
| 6 | GTC seizure with running, jumping, or vocalizations |
| 7 | Tonic full-body extension |
| 8 | Respiratory arrest and death |

Classification criteria adapted from established protocols.(*42*, *43*) This scale was applied continuously during the 20-minute post-injection observation window following pentylenetetrazol (PTZ) challenge across all experimental cohorts. See Methods for details.

**Table S2.** **Antibodies and reagents utilized for flow cytometry, immunofluorescence, and in vivo interventions.**

| **Reagent** | **Clone** | **Dilution** | **Vendor** | **Catalog #** |
| --- | --- | --- | --- | --- |
| ***Flow Cytometry*** | | | | |
| Fc Block (anti-CD16/CD32) | 2.4G2 | 1:100–1:250 | BD Biosciences | 553142 |
| FoxP3–AF488 | FJK-16s | 1:100 | eBioscience | 53-5773-82 |
| GATA-3–PE | TWAJ | 1:100 | eBioscience | 12-9966-41 |
| CD19–PEDazzle594 | 6D5 | 1:400 | BioLegend | 115554 |
| Tbet–PE-Cy7 | 4B10 | 1:100 | BioLegend | 25-5825-80 |
| RORγt–APC | B2D | 1:100 | eBioscience | 17-6981-82 |
| CD3–AF700 | 17A2 | 1:200 | BioLegend | 100216 |
| Thy1.2 (CD90.2)–BV421 | 53-2.1 | 1:400 | BioLegend | 140327 |
| CD11b–BV605 | M1/70 | 1:400 | BD Biosciences | 563015 |
| NK1.1–BV650 | PK136 | 1:400 | BioLegend | 108736 |
| CD4–BV711 | RM4-5 | 1:200 | BioLegend | 100557 |
| CD8α–BV785 | 53-6.7 | 1:200 | BioLegend | 100750 |
| CD45–BUV395 | 30-F11 | 1:400 | BD Biosciences | 564279 |
| TCRγδ–PerCP-Cy5.5 | GL3 | 1:200 | BioLegend | 118118 |
| CD69–FITC | H1.2F3 | 1:100 | Biolegend | 104505 |
| CD62L–PerCP-Cy5.5 | MEL-14 | 1:200 | BioLegend | 104431 |
| CD8α–Pacific Blue | 5H10 | 1:200 | Invitrogen | MCD0828 |
| KLRG1–BV510 | 2F1/KLRG1 | 1:200 | BioLegend | 138421 |
| CD3ε–BV711 | 145-2C11 | 1:200 | BD Biosciences | 563123 |
| CD44–BV785 | IM7 | 1:400 | BioLegend | 103041 |
| CD103–BUV737 | M290 | 1:200 | BD Biosciences | 741739 |
| IFNγ–FITC | XMG1.2 | 1:100 | Biolegend | 505806 |
| IL-17A–PE-Cy7 | TC11-18H10.1 | 1:100 | Biolegend | 506922 |
| IL-5–APC | TRFK5 | 1:100 | BioLegend | 504306 |
| IL-13–eF660 | eBio13A | 1:100 | eBioscience | 50-7133-82 |
| CD3ε–BV510 | 145-2C11 | 1:200 | BioLegend | 100353 |
| CD11b–FITC | M1/70 | 1:400 | BioLegend | 101206 |
| CD369 (Dectin-1/ CLEC7A)–APC | RH1 | 1:200 | BioLegend | 144305 |
| CD64–BV605 | X54-5/7.1 | 1:200 | BioLegend | 139323 |
| CD11c–BV650 | N418 | 1:200 | BioLegend | 117339 |
| CX3CR1–BV785 | SA011F11 | 1:200 | BioLegend | 149029 |
| CD45–FITC | 30-F11 | 1:100 | BioLegend | 103108 |
| CD11b–PE | M1/70 | 1:100 | BioLegend | 101207 |
| Ly6C–APC | HK1.4 | 1:100 | BioLegend | 128015 |
| ***Immunofluorescence*** | | | | |
| Rat anti-GFAP | 2.2B10 | 1:1000 | Invitrogen | 13-0300 |
| Rabbit anti-Iba1 | Polyclonal | 1:1000 | Wako | 019-19741 |
| Guinea Pig anti-Iba1 | Gp311H9 | 1:1000 | Synaptic Systems | 234 308 |
| Chicken anti-NeuN | Polyclonal | 1:500 | Millipore Sigma | ABN91 |
| Guinea Pig anti-NeuN | Polyclonal | 1:500 | Millipore Sigma | ABN90 |
| Rat anti-MHC-II (I-A/I-E) | M5/114.15.2 | 1:500 | eBioscience | 14-5321-82 |
| Rabbit anti-STAT1 | D1K9Y | 1:200 | Cell Signaling Technology | 14994S |
| Syrian hamster anti-CD3ε | 500A2 | 1:200 | BD Biosciences | 553238 |
| Rat anti-CD45–AF488 | 30-F11 | 1:200 | BioLegend | 103122 |
| Mouse anti-Parvalbumin (PV) | PARV-19 | 1:500 | Sigma Aldrich | P3088-100UL |
| Rat anti-CD4–AF647 | RM4-5 | 1:100 | Biolegend | 100530 |
| Rabbit anti-dsRed | Polyclonal | 1:1000 | Takara | 632496 |
| Chicken anti-GFP | Polyclonal | 1:5000 | Aves | GFP-1020 |
| Goat anti-OLIG2 | Polyclonal | 1:200 | R&D Systems | AF2418 |
| Rabbit anti-PKCδ | EPR17075 | 1:2000 | abcam | ab182126 |
| Goat anti-chicken AF488 | Polyclonal | 1:1000 | Invitrogen | A11039 |
| Goat anti-chicken AF555 | Polyclonal | 1:1000 | Invitrogen | A21437 |
| Goat anti-chicken AF647 | Polyclonal | 1:1000 | Invitrogen | A21449 |
| Goat anti-chicken AF405+ | Polyclonal | 1:1000 | Invitrogen | A48260 |
| Goat anti-rabbit AF555 | Polyclonal | 1:1000 | Invitrogen | A21429 |
| Goat anti-rabbit AF647 | Polyclonal | 1:1000 | Invitrogen | A21245 |
| Goat anti-guinea pig AF488 | Polyclonal | 1:1000 | Invitrogen | A11073 |
| Goat anti-guinea pig AF647 | Polyclonal | 1:1000 | Invitrogen | A21450 |
| Goat anti-guinea pig AF405 | Polyclonal | 1:1000 | abcam | ab175678 |
| Goat anti-Syrian hamster AF647 | Polyclonal | 1:1000 | Invitrogen | A21451 |
| Goat anti-rat AF488 | Polyclonal | 1:1000 | Invitrogen | A11006 |
| Goat anti-rat AF555 | Polyclonal | 1:1000 | Invitrogen | A21434 |
| Goat anti-rat AF405+ | Polyclonal | 1:1000 | Invitrogen | A48261 |
| Goat anti-mouse AF488 | Polyclonal | 1:1000 | Invitrogen | A11001 |
| Goat anti-mouse AF555 | Polyclonal | 1:1000 | Invitrogen | A21424 |
| Donkey anti-rabbit AF555 | Polyclonal | 1:1000 | Invitrogen | A31572 |
| Donkey anti-goat AF647 | Polyclonal | 1:1000 | Invitrogen | A21447 |
| Donkey anti-rat AF405+ | Polyclonal | 1:1000 | Invitrogen | A48268 |
| Donkey anti-chicken AF488 | Polyclonal | 1:1000 | Invitrogen | A78948 |
| ***In vivo antibody treatment*** | | | | |
| Rat anti-mouse CD4 | GK1.5 | 250 μg/ mouse | BioXCell | BE0003-1 |
| Rat IgG2b Isotype | LTF-2 | 250 μg/ mouse | BioXCell | BE0090 |

**Table S3.** **Software packages and versions utilized for bioinformatic, statistical, and imaging analyses.**

| **Software** | **Version Used** |
| --- | --- |
| R | 4.2.0 |
| Seurat | 3.4.1 |
| ggplot2 | 3.3.6 |
| dplyr | 1.0.9 |
| Mast | 1.22.0 |
| Matrix | 1.4.1 |
| RStudio | 2022.2.2.485 |
| SigmaPlot | v15.0 |
| OriginPro | v2024 |
| MATLAB | vR2023b |
| Spike2 | 8.21 |
| Graphpad Prism | 11.0.2 |
| FlowJo | v10 |
| Imaris | v9.5.1 |

**Table S4.** **Experimental parameters, quality control thresholds, and clustering metrics for single-cell RNA-sequencing analysis.**

| **Quality Control Parameter / Metric** | **Standard Applied Threshold / Experimental Design** |
| --- | --- |
| Tissue-Specific Cell Yield | 11,863 high-quality microglia (post-filtering) |
| Biological Replicates (mice) | Group 1 (Sham + IgG): 4 mice (2 male, 2 female)  Group 2 (TBI + IgG): 4 mice (2 male, 2 female)  Group 3 (TBI + a-CD4): 4 mice (2 male, 2 female) |
| Microfluidic Processing Architecture | 3 independent 10x Genomics lanes (1 lane pooled per experimental group) |
| Single-Cell Capture Chemistry | Chromium Next GEM Single Cell 3' Reagent Kits v3.1 (Dual Index, Chip G) |
| Unique Molecular Identifier (UMI) Depth | 4,000–20,000 UMIs/cell |
| Feature / Gene Complexity Thresholds | 2000–5000 genes/cell |
| Mitochondrial Transcript Transcriptome Cap | 0–5% mitochondrial RNA/cell |
| Data Normalization & Batch Correction | SCTransform (Seurat package) regressing out nFeature_RNA and percent.mt variance |
| Differential Expression Inclusions | Gene must be productively expressed in ≥5% of cells within the cluster |
| Unsupervised Clustering Resolution | Granularity parameter set to 0.5 via the FindClusters function |
